# STIMscope: centimeter-scale all-optical imaging and patterned optogenetic manipulation at single-cell resolution

**DOI:** 10.64898/2026.05.27.728160

**Authors:** Hamid Chorsi, Saray Soldado-Magraner, Yan Jin, Imaan Soltanalipouryekesammak, Alex Zheng, Dejan Markovic, Daniel H. Geschwind, Peyman Golshani, Dean V. Buonomano, Daniel Aharoni

## Abstract

Linking observation to intervention at cellular resolution makes it possible to move from measuring network activity to testing the contribution of defined neurons or ensembles within the same preparation. All-optical probing provides this capability by combining fluorescence-based readout with targeted optogenetic manipulation. Yet the platforms that deliver this capability remain complex, expensive, and difficult to maintain, requiring specialized expertise that has confined them to a small number of laboratories. They also typically provide fields of view too limited for studies of large, distributed neuronal populations. We address these constraints with the Spatiotemporal Illumination Microscope (STIMscope), a one-photon benchtop platform that integrates large-aperture tandem optics with a small-pixel back-illuminated CMOS sensor, a digital micromirror device for patterned illumination, and a GPU-based processing unit coordinated by a microcontroller for hardware-level synchronization. Ray-tracing simulations and point-spread-function measurements confirm cellular-scale resolution, with imaging lateral FWHM of 5.6 µm at the field center and 5.8 µm at the edge, and excitation lateral FWHM of 5.8 µm at the center and 6.2 µm at the edge, supporting fields of view as large as 14 mm × 11 mm in the demagnified configuration. The accompanying Closed-loop ready Real-time Imaging and Stimulation Pipeline (CRISPI) provides GPU-accelerated calibrated mask projection (26.3 ms latency), online ROI trace extraction, and modular ZeroMQ-based control, with a measured imaging-to-stimulation loop benchmark of 91.6 ms. We validate STIMscope in fixed mouse brain tissue, live iPSC-derived human neuronal cultures, and *ex vivo* organotypic slices of mouse auditory cortex. In organotypic slices, we show that both static and spatiotemporal stimulus identity can be decoded from population activity, revealing reservoir-like population dynamics, and that this decodability remains stable in the same neuronal population over hours. We further show that post-stimulus activity retains information about recent stimuli for several seconds, consistent with short-term memory dynamics. With a bill of materials under $5,000 USD and all mechanical designs, firmware, and software released open-source, STIMscope makes all-optical neuroscience experiments a routine capability accessible to laboratories without specialized optical engineering expertise.

## Introduction

A central challenge in circuit neuroscience is moving beyond observation of neural activity to direct tests of how specific cells and activity patterns contribute to circuit function. All-optical probing approaches meet this challenge by combining fluorescence-based activity readout with optogenetic manipulation, enabling population dynamics to be measured and perturbed within the same preparation^1–11^. Over the past decade, advances in genetically encoded calcium^1,12,13^ and voltage^14,15^ indicators, together with light-sensitive opsins for optogenetic excitation and inhibition^16–21^, have substantially expanded the experimental reach of this approach. These tools have enabled experiments that map functional connectivity^2,4^, perturb candidate circuit mechanisms^22^, and quantify how network activity changes with learning, plasticity, and experience^6,23–25^. Cellular-resolution all-optical experiments have largely been implemented on multiphoton platforms^26–33^, which provide subcellular resolution and deep-tissue access but are expensive, maintenance-intensive, and require substantial expertise to assemble, align, and operate. These constraints concentrate such systems in a small number of specialized laboratories and limit cross-site reproducibility. Expanding the imaging area further compounds these issues, since larger fields of view (FOVs) are typically obtained through tiling, sequential scanning, or multi-plane acquisition, each of which increases acquisition time and reduces effective temporal resolution for experiments that aim to track fast population dynamics across large regions or follow the same networks over long durations.

One-photon all-optical platforms offer a more accessible alternative^10,34–38^, combining widefield imaging with patterned optogenetic stimulation using spatial light modulators such as digital micromirror devices (DMDs) or projector-based illumination, and supporting structured stimulation motifs in both in vitro^35^ and in vivo settings^10,37^. However, current one-photon implementations face tradeoffs that limit either spatial coverage or cellular precision. Systems based on projector- or DMD-coupled widefield illumination^34^ can deliver large fields but do not always preserve robust single-cell resolution across the full imaging area^39,40^, while systems built around high-end scientific complementary metal oxide semiconductor (CMOS) sensors^41,42^ maintain sensitivity over large fields at substantial cost. Large-format scientific cameras often dominate the total instrument budget and remain a major barrier to broader adoption.

A constraint common across these platforms is that they are typically optimized around high-magnification, high-numerical-aperture (NA) objectives that prioritize diffraction-limited performance over FOV coverage. Short focal-length objectives map a small sample area onto the sensor, reducing the imaging area captured per frame, and high-NA objectives are often corrected over a limited FOV, making uniform cellular performance across large areas difficult. STIMscope explores a different design point for this tradeoff: pairing low-magnification, large-aperture optics with a small-pixel CMOS sensor to expand imaging and excitation field coverage while maintaining sufficient sample-plane sampling and optical performance for cellular-scale resolution across the FOV.

A separate but equally limiting gap concerns real-time integration. Many existing platforms deliver stimulation through preprogrammed pattern sequences and analyze the resulting activity offline^43,44^, lacking frame-accurate coordination between mask projection, online trace extraction, and live visualization. As a result, experiments that require tight synchronization between readout and patterned illumination, including closed-loop interventions, activity-triggered stimulation, and event-locked perturbations, remain difficult to implement on most available systems.

We address these constraints with the Spatiotemporal Illumination Microscope (STIMscope), a customizable fluorescence microscope that delivers large-FOV imaging while preserving single-cell resolution and patterned optogenetic control. The core design integrates large-aperture, low-magnification optics with a small-pixel back-illuminated CMOS sensor to expand imaging area while maintaining sufficient detected signal and sample-plane sampling. A DMD provides programmable illumination patterns, and a dedicated synchronization unit distributes hardware triggers that keep image sensor acquisition and pattern delivery aligned at single-frame precision. The accompanying Closed-loop ready Real-time Imaging and Stimulation Pipeline (CRISPI) coordinates acquisition, processing, and projection on a shared hardware timing reference, maintaining frame-accurate alignment between image sensor frames, projected patterns, and logged events. CRISPI uses graphics processing unit (GPU) accelerated calibrated projection for low-latency mask delivery, while supporting online trace extraction, live visualization, and real-time logging of frames, masks, timing events, and extracted signals during experiments.

To validate this platform, we use fixed mouse brain tissue, live iPSC-derived human neuronal cultures, and organotypic slices of mouse auditory cortex. In organotypic *ex vivo* slices, STIMscope enables longitudinal all-optical experiments in which defined optogenetic patterns are projected onto local cortical circuits while calcium activity is recorded from large neuronal populations. This preparation provides a controlled system for testing elementary forms of cortical computation, because the experimenter can define the external input while monitoring the resulting population dynamics over extended timescales^45,46^. We use two paradigms to illustrate this capability. First, we project static bar patterns with different orientations, together with short sequences of these patterns, and show that local cortical population activity can discriminate stimulus identity and retain information about previous stimuli after stimulus offset, showing short-term memory capabilities. Second, we project mirrored leftward and rightward moving bars and demonstrate that the same neuronal population can encode spatiotemporal stimulus direction consistently across repeated recording sessions, as predicted by reservoir computing theory^47^. These experiments demonstrate how STIMscope can be used to probe elementary computations and longitudinal stability of local cortical computations in *ex vivo* networks.

To support reproducibility and broader adoption, all mechanical designs, firmware, and software are released openly, together with installation documentation, reference configurations, and standardized outputs. With a bill of materials under $5,000 USD and a reliance on off-the-shelf components, STIMscope is intended to extend access to state-of-the-art all-optical experiments beyond laboratories with extensive instrumentation resources, and to move these experiments toward routine use in laboratories without extensive optical engineering infrastructure.

## Results

STIMscope is implemented as a benchtop system assembled on a standard optical breadboard (**Fig. 1a**). The instrument comprises a microscope body built around a dual-tandem lens configuration^48,49^, a DMD for patterned illumination, a GPU processing unit, a microcontroller unit (MCU) for hardware synchronization, a CMOS image sensor, and a stage holder with an optional motorized controller, all assembled within an overall envelope of approximately 25 cm × 25 cm × 40 cm and suitable for routine laboratory use. The instrument can be readily configured in either inverted or upright geometries (the upright configuration is shown in Supplementary Fig. 1). A 3D-printed cube houses the dichroic mirror and the adaptor rings used to interface the imaging, excitation, and objective lenses. Additional 3D-printed mounts and adaptors connect the optics to the projection module and align them with the rest of the optical path.

**Fig. 1:**
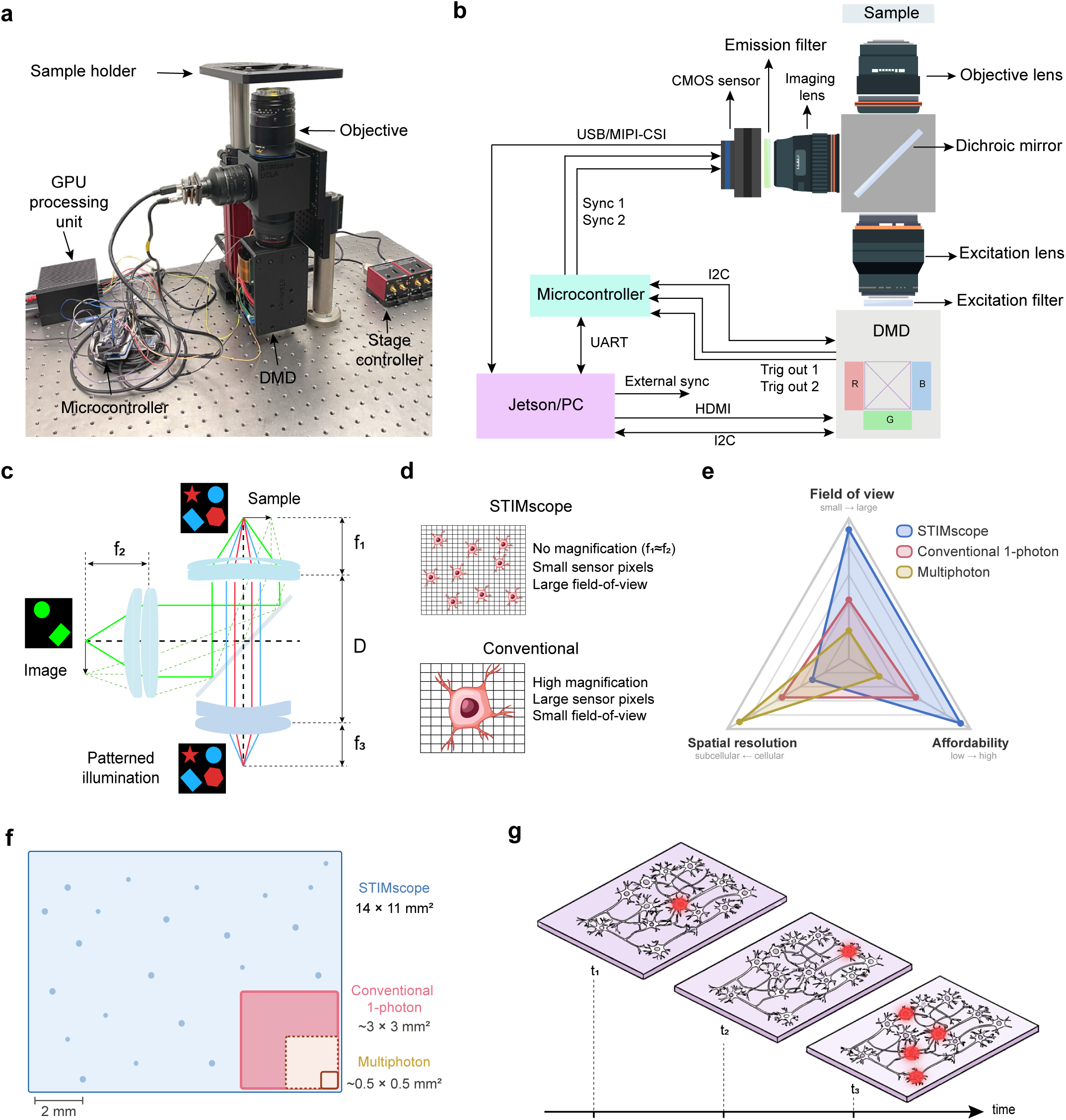
STIMscope platform. **a**, Photo of the implemented STIMscope platform in the inverted configuration. **b**, Hardware architecture for synchronization, control and communication between the image sensor, DMD projector, microcontroller and NVIDIA Jetson Orin in real-time. **c**, Schematic of the optical layout and main components, showing integration of a small pixel CMOS sensor with a low magnification large aperture relay. **d**, Conceptual comparison between conventional high-magnification one-photon imaging and the STIMscope design strategy. **e**, Conceptual comparison of representative STIMscope, conventional one-photon, and multiphoton operating regimes across FOV, affordability, and spatial resolution. The comparison illustrates typical design tradeoffs rather than absolute limits, since conventional one-photon and multiphoton systems span a broad range of implementations. **f**, Representative FOV comparison. Conventional one-photon microscopy is illustrated using typical 10× widefield configurations, where image sensor-limited FOV commonly ranges from approximately 1.3 × 1.3 mm² to 3 × 3 mm² depending on sensor size or adapter magnification. **g**, Schematic examples of DMD-based patterned optogenetic targeting in neuronal samples, showing programmable illumination of individual cells at different positions or multiple spatially distributed cells within the same FOV.

The hardware architecture coordinates four main subsystems: the DMD evaluation module (DLP4710EVM) projector, the CMOS image sensor (Sony IMX334 or IMX290 in the current configuration, although the sensor module is readily replaced by the user’s preferred image sensor), the Microchip ATSAMD51 MCU, housed on the Adafruit Grand Central M4 Express development board, and the NVIDIA Jetson AGX Orin processing unit (**Fig. 1b**). In addition to projecting patterned excitation, the DLP4710EVM provides a global timing reference: its trigger output, which marks the onset of each projected pattern on the DMD, is routed to the MCU for preprocessing and then forwarded to the image sensor to align sensor exposure and readout with projection timing. The DLP4710EVM communicates with the Jetson AGX Orin over a high-definition multimedia interface (HDMI) connection for pattern delivery, and the Jetson configures and monitors the projector using general-purpose input/output (GPIO) lines and an inter-integrated circuit (I2C) control bus. The I2C link enables projector frame-rate configuration, light-emitting diode (LED) intensity and color-mode control, and continuous status readback during experiments. The DMD runs custom firmware that provides control of pattern delivery, including trigger-ON commands, programmable pre- and post-exposure timing, exposure duration, LED drive-current control, and configuration of the DLP4710EVM into external pattern-streaming mode (details in Methods). Two hardware trigger outputs from the DMD (Trig out 1 and Trig out 2) are routed to the MCU, which preprocesses and forwards synchronization signals to the image sensor over dedicated sync lines (Sync 1 and Sync 2). Trig out 1 marks the onset of each frame and is therefore issued once per frame, while Trig out 2 marks the onset of each individual pattern within a frame and can be issued up to 24 times per frame, corresponding to the maximum number of patterns the DMD can deliver per frame (details in Methods), (analysis in Supplementary Fig. 2). The MCU also receives commands from the Jetson over a universal asynchronous receiver/transmitter (UART) serial link and communicates with the DMD over I2C (details in Methods). Image data are transferred from the CMOS sensor, operating in slave mode and synchronized to the projector trigger, to the Jetson over a USB connection in the current configuration, with the option to use a mobile industry processor interface camera serial interface (MIPI-CSI) link on hardware that supports it. Together, this architecture ensures that image sensor exposure and DMD pattern projection are phase-locked to a common hardware timing reference, enabling single-frame synchronization between imaging and patterned illumination.

The optical layout follows a dual-tandem lens design on both the collection and projection paths (**Fig. 1c**). On the imaging path, an objective lens with focal length f₁ collects fluorescence emission from the sample, which is relayed to the CMOS sensor by an imaging lens with focal length f₂, setting the imaging magnification as M = f₂/f₁. On the excitation path, a separate excitation lens with focal length f₃ projects DMD patterns onto the sample plane, with the excitation magnification set independently as M = f₁/f₃. A dichroic mirror (details in Methods and Supplementary Fig. 3) separates the excitation and emission paths, and excitation and emission filters select the appropriate spectral bands. The CMOS sensor must be placed at the flange distance of the imaging lens. Because the DLP4710EVM module places the DMD chip 45 mm from the mechanical entrance aperture, the excitation lens is currently constrained to Nikon F-mount optics (flange focal distance 46.5 mm). Beyond the primary design considerations, the use of large-aperture lenses also provides an additional practical degree of freedom: the working distance can be adjusted by selecting lens mounts with different flange distances (details in Methods).

This architectural choice is compared schematically to a conventional high-magnification microscope (**Fig. 1d**). A conventional system uses high magnification and large sensor pixels to image a small FOV at diffraction-limited resolution. STIMscope instead uses a small-pixel-size image sensor (2 µm pixel pitch for the IMX334) and operates at or near unit magnification (f₁ ≈ f₂), expanding the achievable FOV (centimeter scale, discussion in Supplementary Note 1) while maintaining sufficient sample-plane sampling and optical performance for single-cell imaging and stimulation across the entire FOV (calculations presented in Methods). Rather than relying on larger pixels to cover a wider area, STIMscope tunes the overall magnification to exploit the small pixel pitch, expanding field coverage while preserving cellular precision. This design choice is the central distinction of the STIMscope optical architecture.

These design choices place STIMscope in a distinct operating regime relative to representative conventional one-photon and multiphoton systems (**Fig. 1e**). The comparison is intended to illustrate typical design tradeoffs rather than absolute limits, since both categories span a wide range of implementations. Cellular-resolution multiphoton platforms typically image sub-millimeter to approximately millimeter single-field areas, depending on scan optics, objective field, and off-axis aberrations, and specialized multiphoton mesoscopes^50,51^ can reach larger fields at the cost of substantially more complex optical and scanning architectures. STIMscope occupies a different design space, prioritizing large FOV, affordability, and cellular-scale performance in optically accessible preparations.

The FOV comparison further illustrates this operating point (**Fig. 1f**). STIMscope provides millimeter- to centimeter-scale single-shot imaging, substantially expanding the area captured relative to representative 10× conventional one-photon^52,53^ and standard cellular-resolution multiphoton fields. This larger field reduces the need for tiling or sequential scanning when measuring distributed neuronal activity across large populations.

A central capability of STIMscope is targeted patterned excitation: the setup supports programmable DMD-based illumination patterns across the full FOV with single-cell resolution (**Fig. 1g**). Unlike physical photomasks, aperture wheels, or galvo-scanned spots, where changing the pattern requires swapping or moving hardware, stimulation masks in STIMscope are defined in software and rendered by the DMD, allowing different spatial patterns to be projected onto the same FOV without mechanical movement or optical realignment. This enables targeting of individual cells at different locations, as well as simultaneous illumination and patterned imaging of multiple spatially distributed cells. Together, these hardware, optical, and synchronization features provide the foundation for centimeter-scale fluorescence imaging combined with single-cell targeted patterned optogenetic manipulation.

To evaluate the optical performance of the large-aperture tandem lens configuration, where we define large aperture as operating the lenses at low f-number (f/2.8 to f/5.6, corresponding to entrance pupil diameters on the order of several centimeters depending on focal length), we performed ray-tracing simulations in Zemax OpticStudio across a 12 mm field at two wavelengths of interest, 470 nm and 610 nm (**Fig. 2a**). Spot diagrams computed across the imaging field show root mean square (RMS) spot radii of 2.4 µm and 3.9 µm at the field center (image area, IMA: 0.0 mm), 3.1 µm and 3.4 µm at an intermediate position (IMA: 3.0 mm), and 6.0 µm and 4.1 µm at the field edge (IMA: 6.0 mm), indicating that the tandem large-aperture configuration maintains cellular-scale optical performance across the imaging area (**Fig. 2b**).

**Fig. 2:**
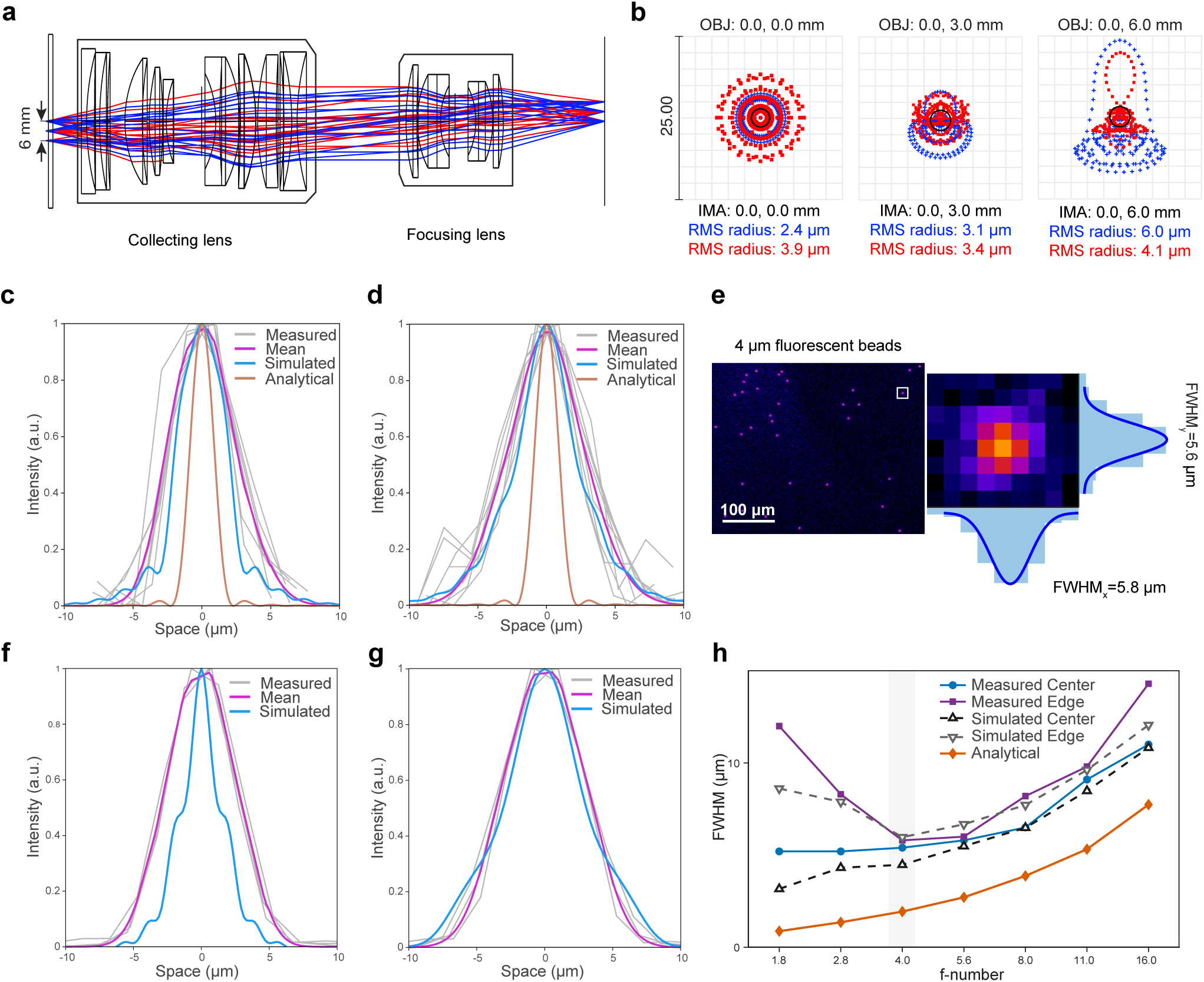
Optical simulation and resolution characterization of STIMscope. **a**, Zemax OpticStudio ray-tracing simulation of the large-aperture tandem-lens configuration used in STIMscope, evaluated at wavelengths relevant to fluorescence imaging (470 nm) and optogenetic excitation (610 nm). **b**, Simulated spot diagrams at field positions 0.0, 3.0, and 6.0 mm from the optical axis. Blue and red traces correspond to the simulated wavelengths, with RMS spot radii reported for each field position. **c**, Imaging PSF measured at the center of the field using 500 nm fluorescent beads. Individual bead profiles are shown in gray, the averaged measured profile is shown in magenta, the Zemax simulation is shown in blue, and the analytical Airy prediction is shown in orange. **d**, Imaging PSF measured at the field edge using 500 nm fluorescent beads, plotted as in c. **e**, Imaging of 4 µm fluorescent microspheres and representative zoomed bead image with orthogonal intensity profiles, showing lateral FWHM values of 5.8 µm in x and 5.6 µm in y. **f**, Excitation PSF measured at the field center by projecting a single DMD pixel onto the sample plane and measuring the resulting illumination spot. **g**, Excitation PSF measured near the field edge, plotted as in f. **h**, Measured and simulated FWHM as a function of lens f-number at the field center and edge, compared with the analytical Airy prediction. The shaded gray region indicates the operating range used for the main characterization, with f/4 providing the best balance between aberration-limited and diffraction-limited performance.

To experimentally validate imaging resolution, we measured the point spread function (PSF) using 500 nm fluorescent beads (details in Methods) at both the center and edge of the FOV, using a 50 mm excitation lens paired with a 50 mm objective lens and a 33 mm imaging lens (**Fig. 2c, d**). Intensity profiles were extracted from isolated beads and averaged across 9 beads and 2 trials. At f/4, the mean lateral full-width at half-maximum (FWHM) was 5.6 µm at the field center and 5.8 µm at the field edge, indicating near-uniform cellular-scale resolution across the full 12 × 7 mm² imaging area. In both cases, measured profiles were compared against Zemax ray-tracing simulations and an analytical diffraction-limited prediction based on the Airy model. The measured PSFs closely followed the simulated profiles.

To further confirm lateral imaging resolution, we imaged 4 µm fluorescent microspheres and extracted intensity profiles along both lateral axes (**Fig. 2e**). The measured FWHM was 5.8 µm along the x-axis and 5.6 µm along the y-axis, confirming consistent cellular-scale imaging performance across both lateral dimensions and agreement with the PSF measurements from 500 nm beads reported above.

The excitation PSF was characterized by projecting a single DMD pixel onto the sample plane and measuring the resulting illumination spot (**Fig. 2f, g**). Across repeated measurements, the mean lateral FWHM of the excitation PSF was 5.8 µm at the field center and 6.2 µm at the field edge, consistent with cellular-scale optogenetic targeting.

Finally, a distinctive feature of STIMscope—enabled by its large-aperture lenses—is the ability to independently tune the effective aperture along both the excitation and emission paths by adjusting the f-number of the corresponding lenses, adding a practical degree of freedom for balancing spatial resolution, field coverage, and signal collection. We characterized PSF performance as a function of lens f-number at both the center and edge of the FOV, comparing measured values against Zemax simulations and the Airy analytical model (**Fig. 2h**). At low f-numbers, increased optical aberrations broadened the PSF beyond the diffraction limit, while at high f-numbers, diffraction broadening and reduced detected signal degraded PSF performance, with the lower signal collection requiring higher analog gain and further raising the effective FWHM. Simulations and measurements showed the same overall trend across the f-number range. The optimal operating point, indicated by the shaded gray region, was found at f/4, providing the best balance between aberration-limited and diffraction-limited regimes at both center and edge positions. The Airy analytical model does not account for lens aberrations and therefore underestimates the measured FWHM at low f-numbers where aberrations dominate. All PSF measurements and characterization data reported in subsequent sections were acquired at this f-number setting.

To demonstrate the large-FOV imaging capability and configurability of STIMscope, we imaged fixed mouse brain slices expressing GCaMP8s across two representative lens configurations (**Fig. 3a**). In the top row, a 50 mm objective lens paired with a 17 mm imaging lens, approximately 3× demagnification, produced a FOV of 14 mm × 11 mm, of which only roughly half is shown because the tissue occupied only part of the available imaging area (see Supplementary Fig. 4 for the full-FOV image). In the bottom row, images were acquired with a 50 mm objective lens and a 50 mm imaging lens, providing unit magnification. Excitation was delivered through the projection path using an 85 mm excitation lens for the demagnified configuration and a 50 mm excitation lens for the unit-magnification configuration. Zoomed insets from the orange- and red-boxed regions show that cellular-scale resolution is preserved in both configurations, with individual neuronal somata clearly resolved at central and peripheral field locations. These results indicate that the large-aperture tandem-lens design maintains cellular resolution across the usable imaging extent in biological tissue and that the system is readily reconfigurable to trade FOV coverage against magnification by swapping lens pairs.

**Fig. 3:**
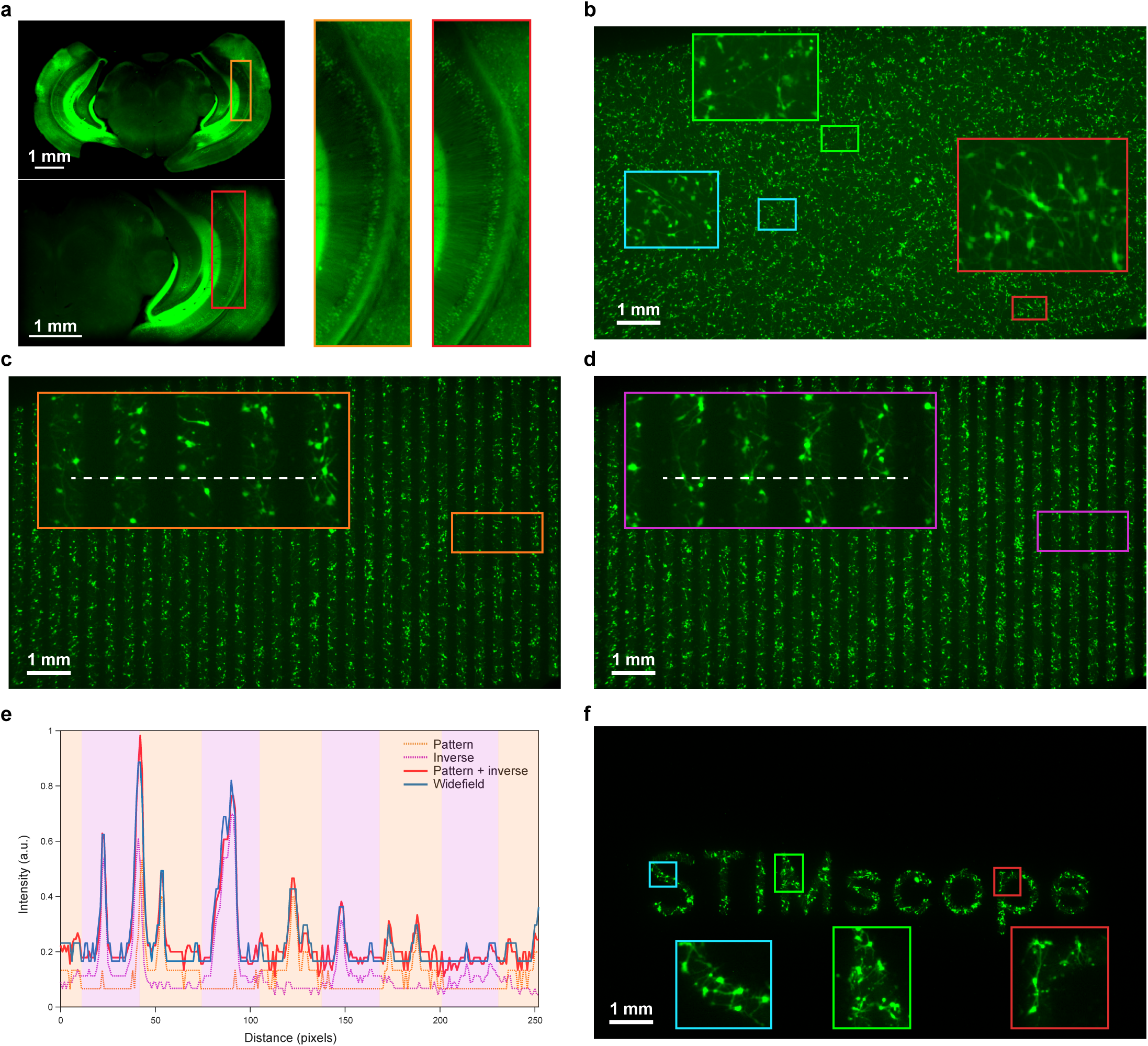
STIMscope imaging and patterned illumination. **a,** Centimeter-scale imaging of fixed mouse brain slices expressing GCaMP8s in two STIMscope configurations. Top, ∼3× demagnification (50 mm objective, 17 mm imaging lens, 85 mm excitation lens; 14 mm × 11 mm FOV; full FOV in Supplementary Fig. 4). Bottom, unit magnification (50 mm objective, 50 mm imaging lens, 50 mm excitation lens). Orange- and red-boxed insets show cellular-scale resolution at the field center and edge. **b,** Widefield reference image of a live iPSC-derived human neuronal culture expressing GCaMP8f in a 24-well plate. Cyan-, green- and magenta-boxed insets confirm single-cell resolution. **c,** Vertical stripe pattern projected across the same culture, with zoomed insets showing selective illumination of neurons in illuminated stripes. **d,** Spatial inverse of the pattern in c projected onto the same culture. **e,** Intensity line profiles along the white dashed lines in c and d for the pattern, its inverse, and their per-pixel sum, compared to the corresponding profile from b. The summed profile matches the widefield reference, indicating high patterning fidelity. **f,** Text pattern "STIMscope" projected across the culture, with zoomed insets showing selective excitation within individual letters.

Having established the imaging performance of the platform in fixed tissue, we next assessed its performance in live preparations by imaging induced pluripotent stem cell (iPSC)-derived human neuronal cultures expressing GCaMP8f (**Fig. 3b–f**). Cultures were grown in 24-well plates with plastic bottoms in standard culture media and maintained in a stage-top incubator throughout imaging to preserve viability and control environmental conditions. Figure 3b shows a representative widefield fluorescence image of the full culture dish, with a 1 mm scale bar illustrating the large-FOV coverage achieved in a single frame. Zoomed insets from the cyan- and magenta-boxed regions at three different scales show that individual neuronal somata and processes remain clearly resolved across the field, indicating that STIMscope maintains cellular-scale resolution in large-FOV live cultures.

We next demonstrated patterned optogenetic illumination in the same live cultures using a series of structured stimulation motifs. Vertical stripe patterns projected across the full culture produced sharp, well-confined illumination bands (**Fig. 3c**), with zoomed insets confirming that neurons within illuminated stripes were selectively illuminated while adjacent neurons in dark stripes remained unilluminated. To show that complementary patterns can be projected onto the same field, we then projected the spatial inverse of this pattern (**Fig. 3d**), producing the expected reciprocal stripe geometry. Horizontal stripe patterns and additional pattern geometries are presented in Supplementary Figs. 5 and 6.

To quantify pattern fidelity, we extracted intensity line profiles along the white dashed lines in Fig. 3c and Fig. 3d and compared the line profiles of the pattern, its spatial inverse, and their per-pixel sum against the corresponding line profile from the widefield reference in Fig. 3b (**Fig. 3e**). If the DMD-defined patterns deliver light to the intended regions, the per-pixel sum of the pattern and its inverse should approximate the widefield profile, because the two complementary masks together span the full illuminated area. The summed profile closely matched the widefield profile across the full line, indicating that the patterned masks deliver light to the intended locations with high spatial fidelity and clear separation between illuminated and dark regions. Small residual deviations between the summed profile and the widefield reference are most likely attributable to ongoing spontaneous activity in the live cultures, slow motion of the culture media, and minor optical scattering from residual debris in the media, rather than systematic patterning errors. The full-FOV relative difference between the widefield image and the summed pattern-plus-inverse image is shown in Supplementary Fig. 7.

Finally, to illustrate the arbitrary-patterning capability of the DMD projection path, we projected the text pattern “STIMscope” across the full culture (**Fig. 3f**). Zoomed insets from the cyan-, green-, and red-boxed regions corresponding to individual letters show that neurons within the illuminated regions were selectively illuminated at cellular-scale resolution, while the sharp boundaries between illuminated and dark regions demonstrate the spatial precision of patterned excitation across complex, user-defined stimulation geometries.

To enable real-time experiments on STIMscope and establish a software foundation for future closed-loop stimulation, we developed the Closed-loop ready Real-time Imaging and Stimulation Pipeline (CRISPI), a modular software system that links synchronized STIMscope hardware to calibrated mask projection, online processing^54,55^, and live visualization (**Fig. 4a**). Building on the hardware synchronization described above, CRISPI provides the software layer that allows time-critical imaging and patterned stimulation experiments to be carried out reliably and reproducibly. Its modular architecture also provides an extensible framework for future closed-loop stimulation strategies that respond to ongoing neural activity. All modules communicate through ZeroMQ^56^ sockets using request/reply (REQ/REP), push/pull, and publish/subscribe (PUB/SUB) messaging patterns, allowing custom components written in any ZeroMQ-compatible language or environment to integrate without modifying the core architecture.

**Fig. 4:**
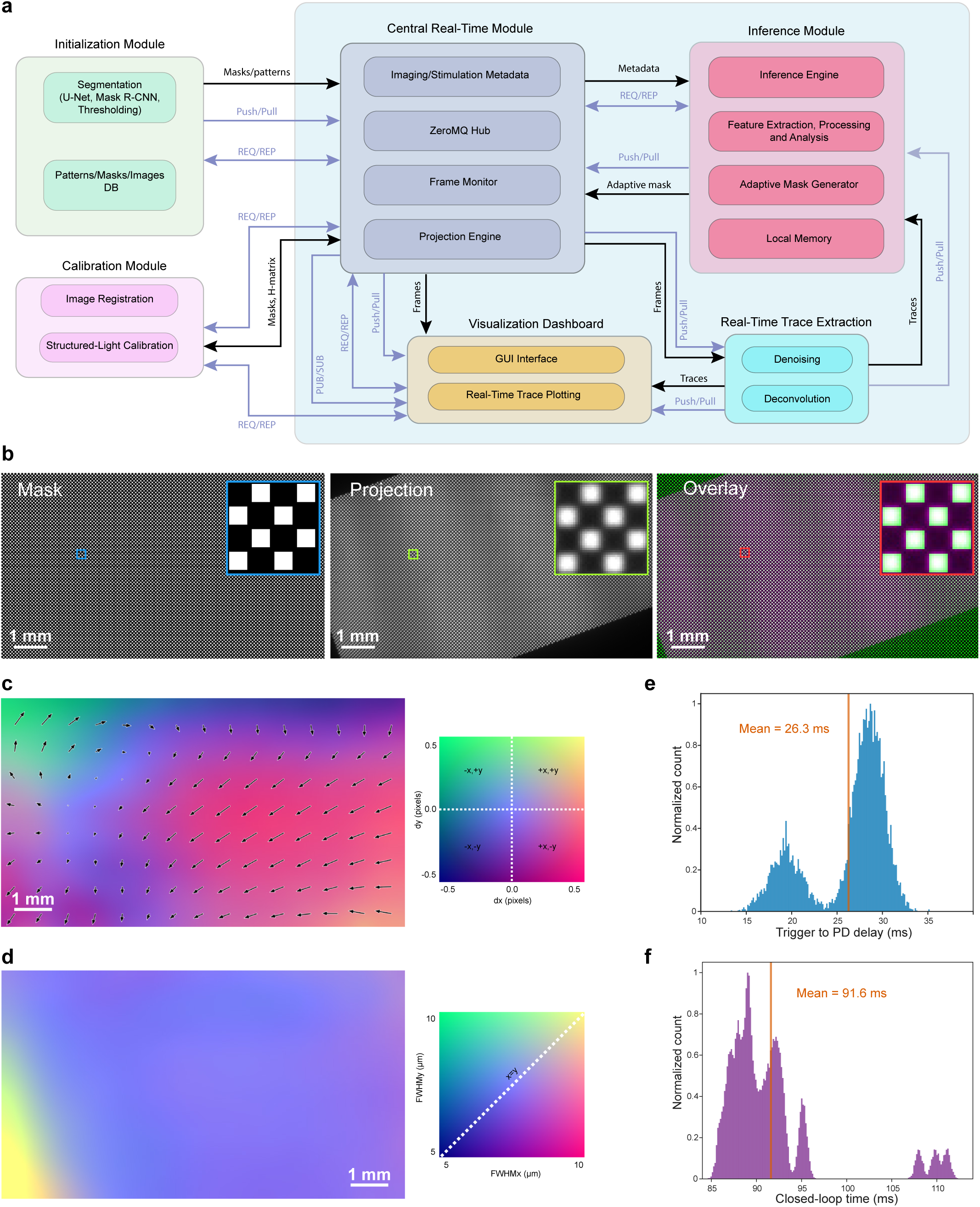
CRISPI architecture, calibrated projection accuracy, and real-time timing performance. **a**, CRISPI software architecture and module-level data flow, comprising initialization, calibration, central real-time coordination, inference, real-time trace extraction, and visualization modules. Modules communicate through ZeroMQ sockets using REQ/REP, PUSH/PULL, and PUB/SUB messaging. Components within the light blue region operate during real-time experiments. **b**, Calibrated mask projection showing the intended square-grid mask (left), captured projection at the sample plane (middle), and direct overlay (right), with the intended mask shown in green and the captured projection shown in magenta. Gray or white regions indicate spatial agreement, while green- or magenta-dominant regions indicate mismatch. **c**, Targeting-error vector field across the image-sensor field, computed from approximately 85,000 single-pixel targets on a 5-pixel grid as measured centroid position minus intended target position. The color map encodes two-dimensional error, and arrows show local error direction and magnitude on a coarse grid. RMS targeting error was 0.46 pixels, approximately 1.3 µm at 2.9 µm per pixel. **d**, Projection PSF width map. FWHMx and FWHMy are encoded as independent color channels, with red indicating broadening along x and cyan indicating broadening along y. Most of the field shows near-isotropic cellular-scale spots, with localized broadening near one corner. **e**, Projection-side latency across 5000 mask presentations, measured using a photodiode at the sample plane. Mean trigger-to-photodiode delay was 26.3 ms. **f**, End-to-end loop timing including mask projection, image-sensor acquisition and transfer, and GPU-based ROI intensity extraction. Mean loop time was 91.6 ms.

CRISPI is organized into six modules: an initialization module, a calibration module, a central real-time module, an inference module, a real-time trace extraction module, and a visualization dashboard (**Fig. 4a**). The modules enclosed within the light blue region operate during real-time experiments. The initialization module runs outside the time-critical loop and handles offline tasks such as neural segmentation using U-Net, Mask R-CNN, or thresholding, along with management of predefined pattern, mask, and image libraries. Its outputs are prepared in advance and passed to the real-time system as needed.

The central real-time module coordinates pattern projection, image acquisition, and online processing relative to the hardware timing reference. Its metadata submodule tracks frame IDs, mask IDs, timestamps, and mask-to-frame correspondence, allowing each projected pattern to be matched to its associated acquired frame. The ZeroMQ hub manages dynamic port binding and message routing between modules. The frame monitor records hardware trigger signals from the projector and the trigger signals forwarded by the MCU to the image sensor, together with associated timing metadata. The projection engine, a major submodule of the central real-time module, receives masks from the initialization, calibration, or inference modules and renders them with low latency for frame-locked delivery. It is implemented as an OpenGL-accelerated GPU renderer that uses texture-transfer optimization, ring buffers, and vsync-gated buffer swaps to minimize latency and ensure deterministic mask delivery.

The calibration module establishes the geometric mapping between the image sensor and the DMD projector, enabling calibrated mask warping prior to display. It supports feature-based calibration using projected registration patterns, such as ChArUco^57^, together with Scale-Invariant Feature Transform (SIFT)^58,59^, Oriented FAST and Rotated BRIEF (ORB)^60,61^, or Affine Scale-Invariant Feature Transform (ASIFT)^62,63^ feature matching to estimate a global 3 × 3 homography matrix. This matrix is passed to the projection engine and applied during GPU rendering so that masks defined in image-sensor coordinates are projected to the corresponding sample locations. For higher-accuracy calibration, the module also supports structured-light calibration using Gray-code^64^ pattern sequences to recover a dense sensor-to-projector lookup table. Details of the calibration and interpolation procedures are provided in Methods.

The inference module serves as the closed-loop extension point in the current CRISPI architecture. It is not implemented in this version of CRISPI, but the modular design defines its interfaces, data flow, and intended role: it is designed to receive imaging metadata and extracted traces, perform feature extraction and analysis, and generate adaptive stimulation masks through an adaptive mask generator. This scaffolding allows future activity-dependent stimulation logic to be developed and integrated without modification to the rest of the pipeline. The real-time trace extraction module receives frames and computes per-region-of-interest (ROI) fluorescence traces, with optional denoising and deconvolution. These traces are routed to the visualization dashboard for live inspection and can also be passed to the inference module for future activity-dependent stimulation workflows. The visualization dashboard provides a graphical user interface with real-time trace plotting and synchronized display of imaging and stimulation events.

We first assessed calibrated mask projection by projecting a square-grid pattern (8 × 8 pixel squares, 10-pixel pitch) after image sensor-to-DMD registration and comparing the intended mask, the captured projection, and their direct overlay (**Fig. 4b**). In the overlay, the intended mask is shown in green and the captured projection in magenta, without thresholding. Regions that appear gray or white indicate spatial agreement between the intended and measured patterns, while green-dominant or magenta-dominant regions indicate mismatch. This provides a direct visual assessment of calibrated projection fidelity across the field of view (further quantification provided in Supplementary Fig. 8).

We then quantified targeting accuracy across the image-sensor field by projecting single-pixel targets on a regular grid with 5-pixel spacing in both axes and measuring the centroid of each captured spot (**Fig. 4c**). The local targeting error was defined as the measured centroid position minus the intended target position. Across approximately 85,000 target locations spanning the 1936 × 1096 pixel image-sensor field, the RMS targeting error was 0.46 pixels, corresponding to approximately 1.3 µm at 2.9 µm per pixel (details in Methods). The composite error map shows sub-pixel targeting accuracy over most of the field, with a smooth residual rotational component and a localized region of larger error near one corner, consistent with residual non-affine distortion not fully captured by a global homography. The continuous error field in Fig. 4c was reconstructed from scattered per-target measurements using the interpolation procedure described in Methods.

We also mapped the projected spot size across the image-sensor field by measuring the FWHM of each captured spot along the x and y axes (details in Methods) (**Fig. 4d**). Across most of the field, the excitation spot remained near-isotropic and cellular-scale, with median FWHM values of approximately 6.1 µm along x and 5.7 µm along y. Spot width increased toward the periphery, with the strongest broadening localized to one corner, consistent with off-axis aberrations such as astigmatism or coma from slight optical decentering or tilt. The majority of the field remained suitable for cellular-scale patterned illumination. As in Fig. 4c, the continuous PSF width map was reconstructed using the interpolation procedure described in Methods. To characterize projection latency, we measured the delay from mask commit to the GPU back buffer to the onset of the photodiode response at the sample plane across 5000 projected masks (details in Methods) (**Fig. 4e**). This measurement captures the projection-side latency, including calibrated mask rendering, GPU buffer management, vsync-gated presentation, HDMI delivery, DMD timing, and LED response. The mean trigger-to-photodiode delay was 26.3 ms. The distribution showed two timing modes, consistent with refresh-period quantization and the timing relationship between mask commit and the next available projector update.

Finally, we measured an end-to-end loop timing benchmark that included mask projection, image sensor acquisition and transfer, and GPU-based processing of all ROIs in a frame by averaging pixel intensities within each ROI (details in Methods) (**Fig. 4f**). This benchmark produced a mean end-to-end loop time of 91.6 ms. The reported timing does not include execution of a future inference model, but it provides a practical measurement of the current real-time pathway from projection through frame capture and ROI-level processing. During acquisition, every frame, mask, trigger, and device event is timestamped and written to structured logs, enabling offline reconstruction of mask-frame correspondence and verification of timing performance. Although developed for STIMscope, CRISPI is designed to be adaptable to imaging systems that expose image sensor frames to the host and accept HDMI- or display-based pattern delivery, including potential applications in patterned illumination and visual-stimulus delivery. Further analyses of projection performance, including projection fidelity and optical crosstalk between adjacent illuminated and dark regions, are provided in Supplementary Figs. 10 and 11.

The preceding sections establish STIMscope as a platform for large-FOV fluorescence imaging with cellular-scale patterned optogenetic illumination, and characterize its optical, projection, and timing performance. In the following sections, we apply the platform to two neuroscience use cases that illustrate how these capabilities translate into experiments addressing concrete circuit-level questions.

To test whether STIMscope can probe elementary computations in local cortical circuits, we performed all-optical experiments in organotypic slices of mouse auditory cortex expressing the calcium indicator GCaMP6f and the red-shifted opsin Chrimson. Patterned optogenetic stimuli were projected onto the cortical surface while population calcium activity was simultaneously recorded from the same field of view. This configuration enabled the delivery of spatially structured inputs while monitoring the resulting activity across large local neuronal populations (**Fig. 5a, b**).

**Fig. 5:**
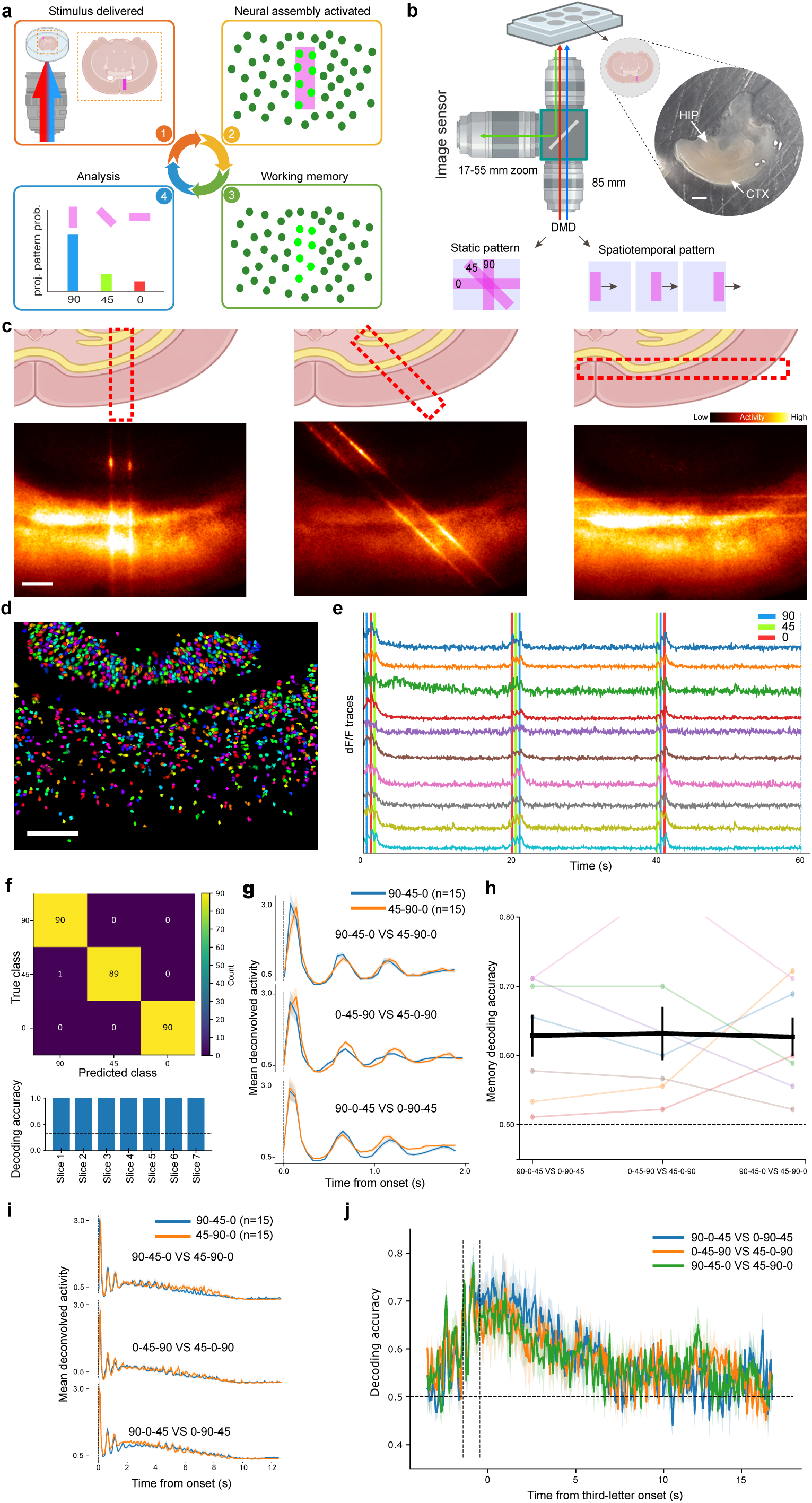
Decoding static patterned stimulation and stimulus short-term memory in organotypic cortical cultures using STIMscope. a, Conceptual schematic illustrating the use of patterned optogenetic stimulation and calcium imaging to probe elementary computations in local cortical circuits. Defined optical patterns are projected onto the culture, activating specific neuronal assemblies, while population activity is recorded for decoding analyses to determine the identity of the projected pattern (e.g., a vertical 90° bar versus 45° and 0° orientations). **b,** Experimental configuration for longitudinal calcium imaging and patterned optogenetic stimulation in organotypic cultures using STIMscope. The platform supports projection of static oriented patterns and spatiotemporal moving-bar stimuli while maintaining the tissue alive in a stage-mounted mini-incubation chamber. Scale bar, 0.5 mm. **c,** Top, example projected static bar patterns at 90°, 45°, and 0° overlaid on organotypic cortical tissue. Bottom, activity elicited on the slice upon pattern projection and optogenetic excitation. Scale bar, 0.5 mm. **d,** Example ROIs extracted from a cortical slice, showing the neuronal population used for decoding analyses. ROIs were identified in both cortical and hippocampal regions. Scale bar, 0.5 mm. **e,** Representative calcium responses from 10 randomly selected ROIs evoked by repeated presentation of the oriented optogenetic patterns. **f,** Population decoding of static stimulus identity. A linear decoder trained on neuronal population activity accurately classifies the three projected orientations. Top, confusion matrix from an individual slice. Each of the six possible triplet permutations was presented 15 times, resulting in 90 total presentations of each individual orientation across the session. Bottom, decoding accuracy across 7 different slices. **g,** Trial-averaged population responses to triplet sequences of oriented patterns using deconvolved calcium activity. Each pattern was presented for 4 frames followed by 4 frames off (15 Hz projection and acquisition rate). **h,** Decoding analysis across pattern sequences, showing that population activity contains information about the recently presented stimulus sequence. **i,** Trial-averaged population responses to triplet sequences of oriented patterns. Persistent post-stimulus activity is observed following sequence presentation, consistent with a short-lived memory trace of recent stimuli. **j,** Time-resolved decoding accuracy for pairwise sequence classification. Decoding remains above chance during and after stimulation for several seconds, indicating that local cortical activity retains information about recent stimulus history.

We first asked whether local cortical population activity could discriminate between static optogenetic patterns. Three bar stimuli of different orientations (90°, 45°, and 0°) were presented in randomly interleaved triplets across trials (e.g., 90°-0°-45°, 0°-45°-90°). These stimuli recruited partially overlapping yet distinct neuronal ensembles, generating reproducible population responses (**Fig. 5c–e**). A linear decoder trained on the evoked population activity during stimulation accurately classified stimulus identity (**Fig. 5f, g**), demonstrating that STIMscope can resolve stimulus-specific cortical population dynamics elicited by different spatial patterns.

We next asked whether cortical population activity retained information about recently presented stimuli after stimulus offset. To test this, we attempted to decode stimulus history from neural activity recorded during triplets sharing the same final stimulus. For example, comparing the sequences 90°-0°-45° and 0°-90°-45°, we asked whether activity during presentation of the final 45° stimulus could reveal which preceding sequence had been delivered. Pairwise decoding between different stimulus sequences revealed above-chance classification performance, consistent with a short-term memory trace embedded in the evoked population dynamics (**Fig. 5h**). Strikingly, decoding accuracy remained elevated not only during presentation of the final stimulus, but also for several seconds after the triplet had ended (**Fig. 5i–j**), indicating that information about recent stimulus history may persist within the recurrent network dynamics. Importantly, all decoding was done using the deconvolved activity, ruling out any memory artifacts generated by the calcium dynamics.

Together, these results demonstrate that STIMscope can be used to study stimulus decoding and short-term memory-like dynamics in ex vivo cortical circuits. By combining patterned optogenetic stimulation with large-field calcium imaging, the platform enables controlled interrogation of how local cortical networks transform defined spatial and temporal inputs into distributed population activity patterns.

We next tested whether STIMscope could support stable decoding of spatiotemporal stimuli across repeated recording sessions. For this experiment, we projected a moving-bar stimulus across the organotypic slice (**Fig. 6a**). The bar moved either leftward or rightward across the same cortical field, generating two mirrored spatiotemporal input patterns. Unlike the previous static-pattern paradigm, this experiment tested whether the temporal structure of population activity evoked by the two directions could be reliably decoded over extended recordings.

**Fig. 6:**
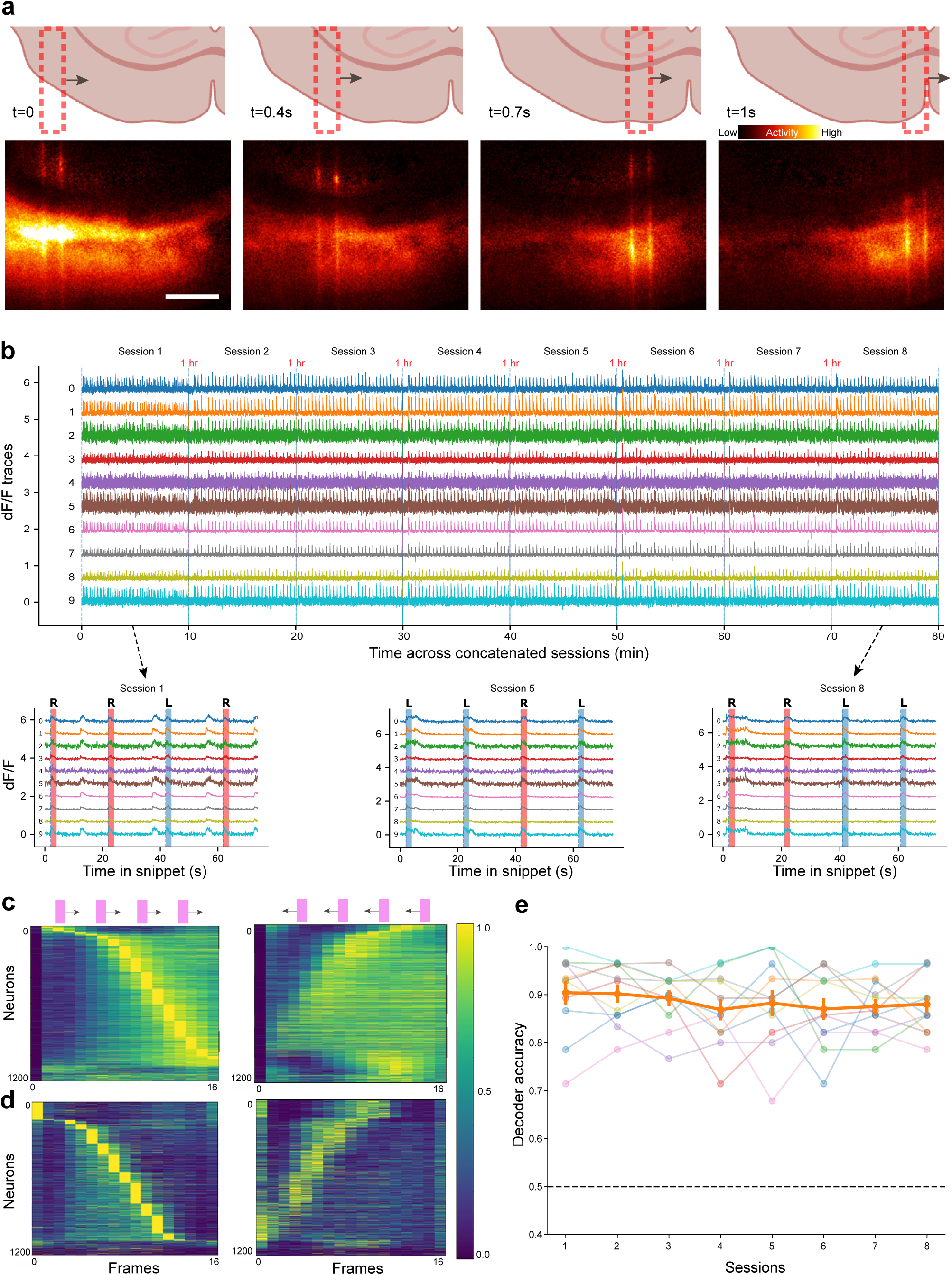
Stable longitudinal decoding of spatiotemporal stimuli in organotypic cortical cultures using STIMscope. a, Top, experimental paradigm for spatiotemporal patterned stimulation. A moving optogenetic bar was projected across the organotypic cortical slice either leftward or rightward, generating two mirrored spatiotemporal input patterns across the same neuronal population. The bar was presented for 1 s at 15 Hz and translated across the field in steps of one bar width. Bottom, example population activity evoked by patterned optogenetic stimulation. Scale bar, 0.5 mm. **b,** Longitudinal calcium imaging of the same neuronal population across eight recording sessions spanning several hours. Example activity traces from 10 randomly selected neurons are shown. Leftward and rightward moving-bar stimuli were repeatedly presented in random order while calcium activity was recorded from the same field of view. Insets show zoomed population responses, demonstrating stable stimulus-locked activity across sessions. In each session, 15 leftward and 15 rightward trials were presented with an inter-stimulus interval of 20 s. Stimulation and recording were performed at 15 Hz. **c,** Trial-averaged calcium responses of the neuronal population during moving-bar stimulation. Neurons were sorted according to the timing of their peak response during rightward-moving trials (left). The same neuronal ordering was preserved for leftward-moving trials (right). **d,** Same analysis as in c using deconvolved neural activity. **e,** Decoding of stimulus direction from population activity across recording sessions. A linear decoder trained on neuronal population responses accurately classified leftward versus rightward moving-bar trials across all eight sessions.

The same neuronal population was imaged across eight recording sessions spanning several hours. During each session, leftward and rightward moving-bar stimuli were presented repeatedly in random order while calcium activity was recorded from the same field of view (**Fig. 6b**). Population responses remained strongly stimulus-locked across sessions, demonstrating stable optical access and reliable evoked activity over time. To visualize the structure of the evoked responses, neurons were sorted according to their response timing (**Fig. 6c**). In the first plot, neurons were ordered by the timing of their peak activity during rightward-moving trials, revealing a sequential activation pattern spanning the stimulus presentation window. The second plot shows responses from the same neuronal population during leftward-moving trials while preserving the ordering obtained from rightward trials. Although the two stimuli are spatial mirror images of one another, the resulting population activity does not exhibit a simple mirrored structure. Instead, the cortical cultures generate distinct distributed activity patterns that discriminate stimulus direction, consistent with reservoir-like dynamics in recurrent cortical circuits^47^. Similar results were observed for the deconvolved activity traces (**Fig. 6d**). We next quantified stimulus discriminability using a linear decoder trained to classify leftward versus rightward trials from population activity. Decoding accuracy remained consistently above chance across all eight sessions, showing that spatiotemporal stimulus direction could be robustly recovered from population responses over hours (**Fig. 6e**). Together, these results demonstrate that STIMscope can longitudinally track the same neuronal population while supporting stable decoding of spatiotemporal inputs in *ex vivo* cortical circuits, revealing rich reservoir-like population dynamics.

## Discussion

STIMscope provides an accessible all-optical platform that combines centimeter-scale FOVs with cellular-resolution fluorescence imaging, patterned optogenetic illumination, and hardware-synchronized real-time control. By pairing low-magnification, large-aperture optics with a small-pixel image sensor and a DMD, the system delivers large-FOV imaging while preserving cellular-scale resolution across the usable field, and the CRISPI pipeline couples this hardware to a synchronized, frame-locked workflow for calibrated mask projection, online trace extraction, and live visualization. With a bill of materials under $5,000 USD and a fully open-source release of mechanical designs, firmware, and software, the platform is intended to move all-optical experiments from a custom engineering effort toward a reproducible method that can be implemented in laboratories without extensive optical engineering infrastructure.

Relative to multiphoton all-optical systems, which provide subcellular resolution and deep-tissue access but are expensive, maintenance-intensive, and concentrated in a small number of specialized laboratories, STIMscope occupies a different design point. It does not aim to provide subcellular resolution, optical sectioning, or imaging in scattering tissue, and is best suited to optically accessible preparations such as cultured neurons and organotypic slices. In exchange, it offers simpler optical design, substantially larger imaging FOVs, and ease of deployment. Imaging and excitation PSF measurements indicate that the tandem-lens architecture preserves cellular-scale resolution across most of the imaging FOV, with the most peripheral corner showing localized broadening consistent with off-axis aberrations from the large-aperture optics. CRISPI addresses the practical challenges of frame-locked operation by combining GPU-accelerated rendering, optimized texture transfers, and vsync-gated buffer swaps to deliver calibrated masks with predictable refresh-quantized timing, with a measured mask-to-light latency of 26.3 ms. A modular ZeroMQ-based architecture and time-stamped logging ensure that acquisition, processing, and projection remain both flexible and fully traceable.

On the projection and acquisition side, mask delivery is currently quantized by the HDMI projector refresh interval, and image data are transferred from the CMOS sensor over USB; both paths can be tightened in future implementations by adopting direct projector-control or pattern-streaming modes in place of HDMI display output, which could reduce or bypass display-refresh quantization, and a MIPI-CSI image sensor interface in place of USB, which could reduce frame-transfer latency. While CRISPI provides a hardware-synchronized framework for online trace extraction and calibrated mask delivery, the inference module that would enable activity-dependent closed-loop stimulation is not implemented in the current version. The modular architecture defines its interfaces, data flow, and intended role, providing a scaffold for future implementations, but full closed-loop stimulation strategies that respond to ongoing neural activity remain to be developed and validated.

The organotypic culture experiments demonstrate how STIMscope can be used to probe local cortical computation in a controlled *ex vivo* preparation. By projecting defined optogenetic patterns while recording calcium activity from large neuronal populations, the platform enables systematic investigation of how cortical circuits encode spatial stimulus identity, temporal stimulus order, and recent stimulation history. The static-pattern experiments show that different oriented inputs evoke distinguishable population responses, while the sequence experiments indicate that cortical activity retains information about recently presented stimuli beyond the stimulation period, consistent with short-timescale memory dynamics.

The moving-bar experiments extend this framework to spatiotemporal stimuli and longitudinal recordings. Mirrored leftward and rightward moving bars evoked distinct population trajectories that could be reliably decoded across repeated sessions spanning several hours. These results show that local cortical networks encode the temporal structure of patterned inputs and that STIMscope can stably track these representations in the same neuronal population over extended experiments. The emergence of distinct distributed population trajectories from mirrored inputs is further consistent with reservoir-like dynamics in recurrent cortical circuits^47^. Together, these results establish STIMscope as a practical platform for studying elementary computations in cortical networks, including stimulus discrimination, short-timescale memory, and longitudinal stability of population dynamics.

More broadly, the combination of centimeter-scale imaging, programmable patterned optogenetic illumination, and frame-locked real-time control that STIMscope and CRISPI provide is intended to enable experiments on circuit dynamics, plasticity, and computation across a range of two-dimensional and slice preparations, including primary cultures, iPSC-derived human neurons, brain organoids, and organotypic slices from multiple cortical and subcortical regions. The open-source release of hardware designs, firmware, and software, together with the modular software architecture, is intended to make these capabilities adaptable: users can swap sensors, optics, filter sets, or stimulation paradigms as their experimental needs evolve, and can extend the CRISPI pipeline by adding modules that integrate over ZeroMQ without modifying the core architecture. We anticipate that this combination of accessibility, modularity, and reproducibility will support both new experimental applications and incremental improvements contributed by the broader community. Several extensions of the platform are natural next steps. On the hardware side, the optical architecture can be reconfigured to support additional indicator and opsin combinations by swapping filter sets and dichroics, and the platform is compatible with alternative image sensors with different pixel sizes and readout rates, allowing the FOV-versus-frame-rate tradeoff to be retuned for specific experiments. On the software side, the inference module provides a defined extension point at which adaptive stimulation algorithms can be added without modifying acquisition or projection, supporting future closed-loop experiments that respond to ongoing neural activity in real time. We expect that contributions from the open-source community will accelerate both hardware and software refinements, broadening the range of preparations and experimental questions to which this class of platform can be applied.

Taken together, STIMscope and CRISPI demonstrate that key capabilities of all-optical neuroscience, namely cellular-scale imaging with targeted patterned optogenetic illumination over network-scale fields, can be delivered by an accessible, reproducible, openly released platform built primarily from off-the-shelf components. By scoping the contribution carefully to the regime in which one-photon, low-magnification, large-aperture optics excel, and by providing synchronized software infrastructure for calibrated projection, online trace extraction, live visualization, and timing verification, STIMscope turns this hardware into a usable experimental method. The platform is intended to broaden participation in real-time all-optical circuit interrogation and to provide a foundation for future closed-loop experiments beyond the laboratories where such systems have historically been concentrated.

## Methods

All experiments were conducted in accordance with National Institutes of Health (NIH) guidelines.

### DLP4710EVM I2C commands

The DLP4710EVM was configured through the DLPC3479 I²C interface. Each transaction consists of an opcode byte followed by command-specific parameter bytes; multi-byte numerical fields are transmitted in little-endian order.

Exposure timing was verified using the Read Validate Exposure Time command, {0x9D, 0x00, 0x03, 0xF8, 0x2A, 0x00, 0x00}. The opcode 0x9D specifies the validation command, 0x00 selects external pattern mode, 0x03 selects the pattern bit-depth and color format used for validation, and the final four bytes encode the requested exposure time as 0x00002AF8 (11,000 µs).

Trigger outputs were configured using the Write Trigger Out Configuration command (opcode 0x92). Trigger Out 1 was enabled with non-inverted polarity and a 0 µs delay using {0x92, 0x02, 0x00, 0x00, 0x00, 0x00}, where 0x02 selects Trigger Out 1 and the four trailing bytes set the delay. Trigger Out 2 was configured identically with {0x92, 0x03, 0x00, 0x00, 0x00, 0x00}, where 0x03 selects Trigger Out 2.

The pattern sequence was configured using the Write Pattern Configuration command, {0x96, 0x01, 0x01, 0x07, 0xF8, 0x2A, 0x00, 0x00, 0x98, 0x08, 0x00, 0x00, 0x88, 0x13, 0x00, 0x00}. The opcode is 0x96; 0x01 selects the pattern format; 0x01 specifies one pattern in the sequence; and 0x07 enables the red, green, and blue illumination channels. The next four bytes set the illumination time as 0x00002AF8 (11,000 µs), the following four bytes set the pre-illumination dark time as 0x00000898 (2,200 µs), and the final four bytes set the post-illumination dark time as 0x00001388 (5,000 µs).

Finally, the system was placed into Light Control, External Pattern Streaming Mode using the Write Operating Mode Select command, {0x05, 0x03}, where 0x05 is the opcode and 0x03 selects external pattern streaming.

### Jetson-microcontroller communication

A custom Python interface on the Jetson transmits data to the MCU over a universal asynchronous receiver/transmitter (UART) link at 9600 bps using the following packet format:

|  |  |  |  |
| --- | --- | --- | --- |
| 0x02 | mode byte | length | data |

The start byte (0x02) marks the byteeginning of each packet and is used by the MCU for packet recognition and framing. The mode byte selects one of three operational modes implemented in the MCU firmware:

Mode A: 60 Hz, 1920 × 1080 resolution, 24 bits per pixel

Mode B: 180 Hz, 8-bit monochrome with 3 patterns per frame

Mode C: 1440 Hz, 1-bit monochrome with 24 patterns per frame

The length byte specifies the number of data bytes that follow, which contain instructions for controlling the DMD module, including setting RGB color, adjusting RGB drive current, and enabling triggers. A schematic of the communication modules and representative oscilloscope traces are provided in Supplementary Fig. 2.

### Dichroic mirror, excitation and emission filters

The STIMscope configuration used for the results reported here employed an excitation filter (Chroma 59003m), a custom dual-band dichroic mirror, and an emission filter (Chroma ET525/50m). The custom dichroic was designed and fabricated by Union Optic, Inc. to meet the optical-quality requirements of the large-aperture relay design (50 mm diameter, 5 mm thickness, surface flatness < λ/4 per 25.4 mm; further design details and transmission spectra are provided in Supplementary Fig. 3). The unit price of the custom dichroic was $500 USD, with bulk orders available at reduced cost. The use of a substrate of this thickness and flatness is important for the STIMscope optical configuration: because the imaging path operates through the dichroic, any deviation from a flat optical surface introduces wavefront distortion that degrades the imaging PSF, particularly in the off-axis regions where the large-aperture rays encounter the dichroic at oblique angles.

An earlier version of the platform used a catalog dual-band dichroic beamsplitter (Chroma 59003bs) with a 1 mm-thick substrate. This dichroic carries a manufacturer-specified surface flatness of 2 waves peak-to-valley per inch, a transmitted wavefront of λ/4 peak-to-valley per inch, parallelism ≤ 5 arc seconds, and a scratch/dig specification of 60/40. While these specifications are appropriate for many standard fluorescence microscopy configurations, in our large-aperture relay the combination of substrate thinness and the resulting surface curvature introduced wavefront aberrations and broadened the imaging PSF, motivating the switch to the thicker, flatter custom dichroic. For users assembling their own STIMscope, a dichroic with substrate thickness and surface-flatness specifications appropriate for low-aberration imaging through a large-aperture optical path is therefore recommended. Filter sets and dichroics can be readily swapped to match different combinations of fluorescent indicators and optogenetic actuators.

### Working distance and aperture tuning

STIMscope supports adjustable working distance by swapping the objective-lens mount on a fixed mechanical assembly. The working distance is set by the flange focal distance of the chosen mount, defined as the mechanical distance from the mount reference surface to the focal plane of the attached lens. In our implementation we used interchangeable photographic mounts, including the Nikon F-mount (flange focal distance 46.5 mm) and the Sony E-mount (18.0 mm). Because each mount places the objective lens at a different distance from the sample, swapping mounts shifts the lens position along the optical axis and changes the working distance without modifying the rest of the optical path. This provides a practical way to accommodate different sample geometries, such as covered well plates, organotypic slice cultures, or thicker sample chambers, using the same imaging and excitation optics.

### Per-pixel photon budget and sample-plane sampling

To compare the design tradeoffs between STIMscope and conventional one-photon microscope configurations, we estimated the expected fluorescence signal per image-sensor pixel rather than total optical light-collection efficiency. This distinction is important because total light collection from a point emitter is governed primarily by numerical aperture (NA) and optical transmission, and high-NA microscope objectives generally collect more emitted light than the lower-NA relay optics used in STIMscope. The relevant design tradeoff for STIMscope is instead the per-pixel photon budget over a large field of view, which depends jointly on NA, system magnification, and image sensor pixel size.

For a uniformly fluorescent sample under matched excitation intensity, exposure time, detector quantum efficiency, and optical transmission, the fluorescence signal recorded by each image-sensor pixel scales approximately with the collected solid angle and the corresponding sample-plane area mapped to that pixel. The collected fluorescence scales with NA^2^, while the sample-plane area represented by one image-sensor pixel scales as (p/M)2, where p is the image sensor pixel size and M is the total magnification. Thus, the expected per-pixel signal can be approximated as:

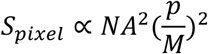

We therefore compared STIMscope with a reference conventional one-photon configuration using the relative per-pixel signal scaling:

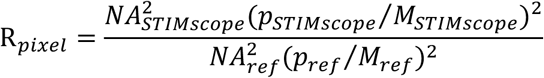

where NA_STIMscope_, p_STIMscope_, and M_STIMscope_ are the numerical aperture, image sensor pixel size, and magnification of STIMscope, and NA_ref_, p_ref_, and M_ref_ are the corresponding values for the reference microscope. A value of R_pixel_ greater than 1 indicates a higher expected fluorescence signal per image-sensor pixel for STIMscope under the stated assumptions, whereas a value of R_pixel_ less than 1 indicates a lower expected fluorescence signal per image-sensor pixel. As an example, using NA_STIMscope_ = 0.3, NA_ref_ = 0.8, p_STIMscope_ = 2 µm, M_STIMscope_ = 1, p_ref_ = 2.9 µm, and M_ref_ = 8: R_pixel = [0.3² × (2/1)²] / [0.8² × (2.9/8)²] = 4.28

This means that, under these assumptions, the expected fluorescence signal per image-sensor pixel is 4.28-fold higher for the STIMscope configuration than for the reference conventional one-photon configuration. This does not mean that STIMscope has higher absolute light-collection efficiency than a high-NA objective. Instead, the increase comes from the larger sample-plane area mapped onto each image-sensor pixel at lower magnification, despite the lower NA.

### PSF measurements

To measure the PSF, we used the TetraSpeck Fluorescent Microspheres Size Kit (Thermo Fisher Scientific), which contains fluorescent microspheres with defined diameters of 0.1, 0.2, 0.5, 1.0, and 4.0 µm, as well as a mixed-size bead region. Microspheres are labeled with four fluorophores with excitation/emission peaks at 365/430 nm, 505/515 nm, 560/580 nm, and 660/680 nm. Beads were imaged directly on the slide under the same optical configuration used for STIMscope characterization. For lateral PSF estimation, isolated 500 nm beads were selected from both the center and edge of the FOV. Intensity profiles were extracted through the bead centers, background-subtracted, normalized, and averaged across beads and trials. The lateral FWHM was then calculated from the averaged intensity profiles.

### Human iPSC culture

Human iPSCs (KOLF2.2J) were maintained in mTeSR Plus (STEMCELL #100-276). Cells were dissociated with Accutase (STEMCELL #07920). Dishes were pre-coated with Geltrex (1:100; Gibco #A1413201) and plated at 50,000 cells/cm2.

#### Lentivirus production

Lentivirus were generated by transfecting HEK293T cells with lentivirus packaging plasmids as previously described^65^ using Lipofectamine 3000 (L3000015): pMDLg/pRRE (Addgene 12251), pRSV-Rev (Addgene 12253) and pCMV-VSV-G (Addgene 8454). To generate GCaMP8f-expressing lentivirus, plasmid FSW-jGCaMP8f (addgene, 197034) was used.

#### Induction of human neurons from human iPSCs

Human iPSCs (either Ngn-2 or Ascl1/Dlx2 integrated at AAV locus) were dissociated and plated at 75,000 cells/cm2 in mTeSR plus Media supplemented with Y-27632. After 72 hours in Neurobasal Media, neurons were dissociated with Accutase and replated either 80,000 /cm2 (Ngn-2 Neurons) or 20,000/cm2 (Ascl1/Dlx2 Neurons). Mouse astrocytes were co-plated with neurons at 120,000/cm2.

### Pixel-to-pixel mapping and calibration

CRISPI uses two complementary calibration strategies: (a) a feature-based homography for rapid global alignment, and (b) a structured-light routine that produces dense, sub-pixel image-sensor-to-projector correspondences for higher-accuracy warping.

To estimate a 3 × 3 homography H_(s → p)_ mapping image-sensor pixels to projector pixels, we project a registration image at native projector resolution and capture the corresponding frame on the image sensor. Prior to matching, both the reference image and the captured image are contrast-normalized using histogram equalization to improve feature detectability under uneven illumination. Local image features are then detected and matched using Scale-Invariant Feature Transform (SIFT), which produces keypoints and descriptors that are robust to scale and rotation changes. Descriptor matching is performed using a two-stage strategy: we first attempt brute-force matching with cross-checking for high-confidence correspondences, and if the number of matches is insufficient, we switch to k-nearest-neighbor (KNN) matching and reject ambiguous matches using the Lowe ratio test (ratio threshold 0.65). The homography is fit using Random Sample Consensus (RANSAC) with a 3-pixel reprojection threshold to exclude outlier correspondences. If RANSAC yields an unstable solution, for example due to low inlier counts, we fall back to Least Median of Squares (LMedS) regression and, if needed, a least-squares refinement.

For diagnostic purposes, the pipeline generates a warped preview image and stores the homography parameters to disk (homography_sensor2proj.npy). In addition to SIFT, the pipeline also supports Oriented FAST and Rotated BRIEF (ORB) and Affine SIFT (ASIFT) feature detectors, as well as ChArUco-based registration patterns, which can be selected at calibration time. Under the planar-scene assumption, the homography corrects the full planar projective transformation between the projector and image-sensor views, including translation, in-plane rotation, uniform and non-uniform scale, shear, and perspective skew (keystone). It does not correct lens distortion or depth-varying warps, motivating the dense structured-light calibration described below.

For sub-pixel alignment and to compensate for spatially varying distortions that are not well captured by a single homography, such as lens distortion, we use a structured-light calibration routine. The projector displays horizontal and vertical Gray-code sequences, which encode projector pixel coordinates as binary patterns that are decoded at each image-sensor pixel to produce dense forward maps from image-sensor coordinates to projector coordinates. The resulting dense lookup table is passed to the projection engine and used to resample stimulation masks at projection time.

### Projection engine

The projection engine is implemented in C++ and uses OpenGL with GLFW for real-time rendering, GLEW for OpenGL extension loading, ZeroMQ for interprocess communication, and libgpiod for hardware trigger input and output through general-purpose input/output (GPIO) lines. Masks and associated metadata are streamed to the projection engine through ZeroMQ sockets. Mask images are received through a PUSH/PULL channel, control messages such as updated calibration transforms are handled through a request/reply (REQ/REP) channel, and status messages such as projected-frame and visibility events are broadcast through a publish/subscribe (PUB/SUB) channel. Rendering is accelerated on the GPU. Incoming masks are uploaded as OpenGL textures and rendered to the projector output. To reduce latency and avoid blocking during texture transfer, the engine uses pixel buffer objects (PBOs), ping-pong or triple buffering, and GPU synchronization fences. The rendering path supports immediate mask scheduling and vsync-gated buffer swaps, allowing masks to be delivered with low latency while remaining synchronized to the projector refresh cycle. Frame-accurate synchronization is achieved by monitoring projector and image-sensor trigger lines using libgpiod. On each projector trigger, the engine increments a projector frame index, determines which mask became visible on that projector frame, and publishes a JSON status message containing the projector index and visible mask identifier. These status messages are used by the graphical interface and downstream modules to maintain deterministic correspondence between projected masks, acquired image frames, and logged experimental events.

### Real-time trace extraction

The visualization dashboard performs real-time trace extraction from the incoming image-sensor stream while maintaining a continuously updated live preview. Frames are handled as NumPy arrays within the PyQt5-based interface. For each incoming frame, the module computes per-ROI fluorescence summaries, such as mean pixel intensity within each ROI, and appends these values to continuously updated traces. Live traces are displayed alongside the image preview using PyQtGraph, which supports high-refresh-rate plotting with low rendering latency. Each trace sample is timestamped and associated with the corresponding image frame, projected mask, and projector event when available through the CRISPI status channels. Projector timing and visibility events are received over ZeroMQ, allowing extracted traces to be aligned with stimulation timing at frame-level accuracy. During each experiment, extracted traces and associated metadata, including ROI definitions, timestamps, sampling rate, calibration identifiers, session configuration, and software version information, are written to disk in machine-readable formats such as NumPy or CSV files with JSON metadata. This structure supports downstream analysis, verification of mask-frame correspondence, and reproducible reconstruction of each experiment.

### Calibration, targeting accuracy, and projection PSF mapping

To quantify spatial targeting accuracy, projected target locations were defined in image-sensor coordinates and converted to DMD coordinates using the calibrated sensor-to-projector transform. Single-pixel projection targets were presented across the accessible field on a regular grid with 5-pixel spacing in both axes. For each projected target, the corresponding captured image was recorded, the spot centroid was estimated, and the local targeting error was computed as (Δx, Δy) = (x_measured_ − x_target_, y_measured_ − y_target_). Approximately 85,000 target locations were sampled across the 1936 × 1096 pixel accessible field, corresponding to 5614 × 3178 µm at a 2.9 µm pixel pitch. The RMS targeting error was computed across all valid target locations. Projection PSF width was quantified from the same spatial sampling procedure. For each captured spot, intensity profiles were extracted along the x and y axes through the measured centroid, and the FWHM was computed independently for each direction. The resulting per-target measurements produced scattered samples of Δx, Δy, FWHMx, and FWHMy across the accessible field.

### Interpolation of targeting-error and PSF width maps

The per-target targeting-error and PSF width measurements were acquired across the accessible field on a regular grid with 5-pixel spacing in both axes. For visualization in Fig. 4c and Fig. 4d, per-target measurements were averaged within coarse spatial bins to define a regular field at reduced resolution, which was then upsampled to pixel resolution for display using tensor-product cubic B-spline interpolation (scipy.interpolate.RectBivariateSpline, spline order k_x = k_y = 3). The quantitative results reported in the main text (RMS targeting error and median FWHM values) were computed directly from the per-target measurements and do not depend on the interpolation procedure.

### Projection and loop timing measurements

Projection latency was measured by placing a photodiode (BPW34 Silicon PIN) at the sample plane and recording the delay between a GPIO timestamp generated when the mask was committed to the GPU back buffer and the rise of the photodiode signal. A total of 5000 masks were projected, and the trigger-to-photodiode delay was computed for each trial. This latency includes GPU rendering, calibrated mask warping, buffer management, vsync-gated presentation, HDMI transfer, DMD timing, and LED response. The DMD was configured through I2C commands to use pattern projection mode, external pattern streaming, defined exposure and pre/post-exposure dark intervals, and LED current settings. End-to-end loop timing was measured by projecting a mask, capturing the corresponding frame on the image sensor, transferring the frame to the host, and computing ROI fluorescence values on the GPU by averaging pixel intensities within all ROIs in the frame. The reported loop time therefore includes projection, image acquisition, frame transfer, and GPU-based ROI extraction, but does not include inference model execution or adaptive mask selection.

### Organotypic culture preparation

Cortical organotypic slice cultures were obtained from postnatal day 5-7 wild-type FVB mice of either sex. Organotypic cultures were prepared using the interface method^46^. Coronal slices (400 µm thickness) containing primary auditory and somatosensory cortex were sectioned using a vibratome (Leica VT1200) and bisected before being individually placed onto Millicell cell culture inserts (MilliporeSigma) in a 6-well plate with 1 mL of culture media per well. Culture media was changed at 1 and 24 h after initial plating and every 2–3 days thereafter. Cutting media consisted of MEM (Corning 15-010-CV) plus (final concentration in mM): MgCl2, 3; glucose, 10; HEPES, 25; and Tris-base, 10. Culture media consisted of MEM (Corning 15-010-CV) plus (final concentration in mM): glutamine, 1; CaCl2, 2.6; MgSO4, 2.6; glucose, 30; HEPES, 30; ascorbic acid, 0.5; 20% horse serum, 10 units/L penicillin, and 10 μg/L streptomycin. Slices were incubated in 5% CO2 at 35 °C. At day-in-vitro (DIV) 5, slices were transfected with the viruses pAAV-hSyn-DIO-ChrimsonR-mRuby2-ST (AAV9) + pENN.AAV.CamKII 0.4.Cre.SV40 (AAV9), at a final concentration of 10^11^ (resulting in a sparse Chrimson transfection) and a full GCaMP transfection with the virus pAAV-CAG-SomaGCaMP6f2 (AAV9) at a final concentration of 10^13^. Each slice received a total of 1 µL of viral cocktail gently delivered via a sterilized pipette above the cortex. Recordings were performed between DIV 22-30 to allow sufficient time for viral expression.

### Tissue imaging and patterned optogenetic stimulation

Organotypic cortical slices were imaged and stimulated using the STIMscope. Slices were transferred to a miniaturized incubation chamber (UNO-T-H-CO2, Okolab) mounted on the microscope stage and continuously maintained at 35 °C and 5% CO₂ throughout the experiment. Imaging was performed using the scope’s CMOS image sensor at frame rates of 15Hz, allowing longitudinal recordings over multiple hours.

Excitatory neurons co-expressing SomaGCaMP6f2 and ChrimsonR were stimulated using the STIMscope to project static and spatiotemporal light patterns onto the cortical surface. Static stimuli consisted of oriented bar patterns at 90°, 45°, and 0°. Spatiotemporal stimuli consisted of a moving 90° bar traversing the cortical field either leftward or rightward. The moving bar was presented for 1 s at 15 Hz and translated across the field in steps of one bar width. Patterned stimulation was delivered using combined red and blue illumination to enable optogenetic excitation while preserving calcium imaging.

### Static-pattern and sequence experiments

For static-pattern decoding, oriented bar stimuli were repeatedly projected onto the cortical culture while calcium activity was recorded from the neuronal population. Sequence experiments consisted of all six permutations of three oriented patterns (90°, 45°, and 0°). Each sequence was presented 15 times in random order with a 20 s interstimulus interval. Individual patterns were presented for 4 frames followed by 4 frames off at 15 Hz acquisition. Population activity during and after stimulation was analyzed to determine whether cortical dynamics retained information about recent stimulus history.

### Longitudinal moving-bar experiments

For spatiotemporal decoding, leftward and rightward moving-bar stimuli were repeatedly presented across eight recording sessions spanning several hours. In each session, 15 leftward and 15 rightward trials were presented in random order with a 20 s interstimulus interval. The same field of view and ROI population were maintained across sessions, allowing longitudinal tracking of the same neuronal population over time.

### Calcium imaging analysis

Calcium imaging data were motion-corrected and segmented using Suite2p. ROIs corresponding to putative neuronal somata were extracted, and fluorescence traces were converted to ΔF/F relative to baseline activity. For all decoding, fluorescence traces were deconvolved to estimate event-related neural activity. For the deconvolution, we used the CaLab package^66^. Trials were aligned to stimulus onset, and population response matrices were constructed from the activity of all detected ROIs.

### Decoding analysis

Stimulus identity was decoded from neuronal population activity using linear support vector machine classifiers implemented with scikit-learn. For static-pattern experiments, classifiers were trained to discriminate 90°, 45°, and 0° stimuli. For sequence experiments, classifiers were trained on pairwise comparisons between sequence identities. For moving-bar experiments, classifiers were trained to discriminate leftward versus rightward trials. Decoder performance was evaluated using leave-one-out cross-validation within each recording session. Time-resolved decoding was computed by training and testing classifiers on population activity in frame-based time windows aligned to stimulus onset.

## Acknowledgements

We are grateful to Dr. Yang Dan and Dr. Lihui Lu (University of California, Berkeley) and Dr. Sattar Khoshkhoo (Brigham and Women’s Hospital) for testing the first STIMscope prototype, validating the platform, and providing valuable feedback throughout its development. We thank Dr. Long Yang for providing the fixed mouse brain slice samples used for imaging, and Federico Sangiuliano Jimka for assistance with assembling the STIMscope setup used for the organotypic mouse slice experiments.

Funding Sources: NINDS U01NS126050 (D.A.), NIMH DP2MH129986 (D.A.), Ressler Neuroscience Translation Award (D.A.), NIDA R01DA060229 (D.V.B)

## Author contributions

H.C. and D.A. conceived the project and overall imaging and stimulation design. H.C. designed, fabricated, assembled, and characterized the STIMscope platform, with input and mentorship from D.A. Y.J. cultured the iPSC-derived human neurons under the supervision of P.G. S.S. performed the organotypic mouse slice experiments under the supervision of D.V.B. H.C. and S.S. jointly assembled the STIMscope setup used for the organotypic experiments. H.C. and S.S. analyzed the data with input from D.A. and D.V.B. H.C. wrote the initial manuscript draft with input from all authors. All authors reviewed and approved the final manuscript.

## Data availability

All design files for constructing STIMscope and implementing CRISPI are publicly available through the Aharoni Lab GitHub repository (https://github.com/Aharoni-Lab/STIMscope). The repository includes hardware schematics and mechanical design files, optical simulation data, and example datasets (including recording videos and calcium imaging data), along with the full CRISPI software codebase and installation and usage documentation.

## References

1. Lin, M. Z. & Schnitzer, M. J. Genetically encoded indicators of neuronal activity. Nat Neurosci 19, 1142–1153 (2016).

2. Spampinato, G. L. B. et al. All-optical inter-layers functional connectivity investigation in the mouse retina. Cell Reports Methods 2, 100268 (2022).

3. Grimm, C. & Emiliani, V. All-optical neurophysiology with holographic light shaping. Photoniques 26–29 (2025) doi:10.1051/photon/202513426.

4. Chen, I.-W. et al. High-throughput synaptic connectivity mapping using in vivo two-photon holographic optogenetics and compressive sensing. Nat Neurosci 28, 2141–2153 (2025).

5. Emiliani, V., Cohen, A. E., Deisseroth, K. & Häusser, M. All-Optical Interrogation of Neural Circuits. J. Neurosci. 35, 13917–13926 (2015).

6. Carolan, J. et al. All-optical voltage interrogation for probing synaptic plasticity in vivo. Nat Commun 16, 8834 (2025).

7. Zhang, Z. et al. A real-time all-optical interface for dynamic perturbation of neural activity during behavior. Cell Reports Methods 5, 101180 (2025).

8. Papaioannou, S. & Medini, P. Advantages, Pitfalls, and Developments of All Optical Interrogation Strategies of Microcircuits in vivo. Front. Neurosci. 16, 859803 (2022).

9. Zhang, Z., Russell, L. E., Packer, A. M., Gauld, O. M. & Häusser, M. Closed-loop all-optical interrogation of neural circuits in vivo. Nat Methods 15, 1037–1040 (2018).

10. Mulholland, H. N., Jayakumar, H., Farinella, D. M. & Smith, G. B. All-optical interrogation of millimeter-scale networks and application to developing ferret cortex. Journal of Neuroscience Methods 403, 110051 (2024).

11. Han, J. L., Heinson, Y. W., Chua, C. J., Liu, W. & Entcheva, E. CRISPRi gene modulation and all-optical electrophysiology in post-differentiated human iPSC-cardiomyocytes. Commun Biol 6, 1236 (2023).

12. Sakamoto, M. & Yokoyama, T. Probing neuronal activity with genetically encoded calcium and voltage fluorescent indicators. Neuroscience Research 215, 56–63 (2025).

13. Zhang, Y. & Looger, L. L. Fast and sensitive GCaMP calcium indicators for neuronal imaging. The Journal of Physiology 602, 1595–1604 (2024).

14. Xu, Y., Zou, P. & Cohen, A. E. Voltage imaging with genetically encoded indicators. Current Opinion in Chemical Biology 39, 1–10 (2017).

15. St-Pierre, F., Chavarha, M. & Lin, M. Z. Designs and sensing mechanisms of genetically encoded fluorescent voltage indicators. Current Opinion in Chemical Biology 27, 31–38 (2015).

16. Deisseroth, K. Optogenetics: 10 years of microbial opsins in neuroscience. Nat Neurosci 18, 1213–1225 (2015).

17. Emiliani, V. et al. Optogenetics for light control of biological systems. Nat Rev Methods Primers 2, 55 (2022).

18. Fenno, L., Yizhar, O. & Deisserothp, K. The Development and Application of Optogenetics. Annu. Rev. Neurosci. 34, 389–412 (2011).

19. Bansal, A., Shikha, S. & Zhang, Y. Towards translational optogenetics. Nat. Biomed. Eng 7, 349–369 (2022).

20. Wang, T., Nonomura, T., Lan, T.-H. & Zhou, Y. Optogenetic engineering for ion channel modulation. Current Opinion in Chemical Biology 85, 102569 (2025).

21. Lorca-Cámara, A., Blot, F. G. C. & Accanto, N. Recent advances in light patterned optogenetic photostimulation in freely moving mice. Neurophoton. 11, (2024).

22. Russell, L. E. et al. All-optical interrogation of neural circuits in behaving mice. Nat Protoc 17, 1579–1620 (2022).

23. Hochbaum, D. R. et al. All-optical electrophysiology in mammalian neurons using engineered microbial rhodopsins. Nat Methods 11, 825–833 (2014).

24. Fan, L. Z. et al. All-Optical Electrophysiology Reveals the Role of Lateral Inhibition in Sensory Processing in Cortical Layer 1. Cell 180, 521–535.e18 (2020).

25. Werley, C. A., et al. All-Optical Electrophysiology for Disease Modeling and Pharmacological Characterization of Neurons. CP Pharmacology 78, (2017).

26. Adesnik, H. & Abdeladim, L. Probing neural codes with two-photon holographic optogenetics. Nat Neurosci 24, 1356–1366 (2021).

27. Helmchen, F. & Denk, W. Deep tissue two-photon microscopy. Nat Methods 2, 932–940 (2005).

28. Papagiakoumou, E., Ronzitti, E. & Emiliani, V. Scanless two-photon excitation with temporal focusing. Nat Methods 17, 571–581 (2020).

29. Triplett, M. A. et al. Rapid learning of neural circuitry from holographic ensemble stimulation enabled by model-based compressed sensing. Nat Neurosci 28, 2154–2165 (2025).

30. Chaigneau, E. et al. Two-Photon Holographic Stimulation of ReaChR. Front. Cell. Neurosci. 10, (2016).

31. Szczypkowski, P., Pawlowska, M. & Lapkiewicz, R. 3D super-resolution optical fluctuation imaging with temporal focusing two-photon excitation. Biomed. Opt. Express 15, 4381 (2024).

32. Pégard, N. C. et al. Three-dimensional scanless holographic optogenetics with temporal focusing (3D-SHOT). Nat Commun 8, 1228 (2017).

33. Adam, Y. et al. Voltage imaging and optogenetics reveal behaviour-dependent changes in hippocampal dynamics. Nature 569, 413–417 (2019).

34. Bhatia, A., Moza, S. & Bhalla, U. S. Patterned Optogenetic Stimulation Using a DMD Projector. in Channelrhodopsin (ed. Dempski, R. E.) vol. 2191 173–188 (Springer US, New York, NY, 2021).

35. Farhi, S. L. et al. Wide-Area All-Optical Neurophysiology in Acute Brain Slices. J. Neurosci. 39, 4889–4908 (2019).

36. Xie, H. et al. Multifocal fluorescence video-rate imaging of centimetre-wide arbitrarily shaped brain surfaces at micrometric resolution. *Nat*. Biomed. Eng 8, 740–753 (2023).

37. Yoshida, R. et al. High-resolution optogenetics generates distinguishable neocortical activity patterns in awake mice. Neuroscience Research 223, 105012 (2026).

38. Zhang, J. et al. A one-photon endoscope for simultaneous patterned optogenetic stimulation and calcium imaging in freely behaving mice. *Nat*. Biomed. Eng 7, 499–510 (2022).

39. Cardin, J. A., Crair, M. C. & Higley, M. J. Mesoscopic Imaging: Shining a Wide Light on Large-Scale Neural Dynamics. Neuron 108, 33–43 (2020).

40. Vanni, M. P., Chan, A. W., Balbi, M., Silasi, G. & Murphy, T. H. Mesoscale Mapping of Mouse Cortex Reveals Frequency-Dependent Cycling between Distinct Macroscale Functional Modules. J. Neurosci. 37, 7513–7533 (2017).

41. Shi, R. et al. Random-access wide-field mesoscopy for centimetre-scale imaging of biodynamics with subcellular resolution. Nat. Photon. 18, 721–730 (2024).

42. Doran, P. R. et al. Widefield in vivo imaging system with two fluorescence and two reflectance channels, a single sCMOS detector, and shielded illumination. Neurophoton. 11, (2024).

43. Sheng, W., Zhao, X., Huang, X. & Yang, Y. Real-Time Image Processing Toolbox for All-Optical Closed-Loop Control of Neuronal Activities. Front. Cell. Neurosci. 16, 917713 (2022).

44. Bowen, Z. et al. NeuroART: Real-Time Analysis and Targeting of Neuronal Population Activity during Calcium Imaging for Informed Closed-Loop Experiments. eNeuro 11, ENEURO.0079-24.2024 (2024).

45. Goel, A. & Buonomano, D. V. Temporal Interval Learning in Cortical Cultures Is Encoded in Intrinsic Network Dynamics. Neuron 91, 320–327 (2016).

46. Stoppini, L., Buchs, P.-A. & Muller, D. A simple method for organotypic cultures of nervous tissue. Journal of Neuroscience Methods 37, 173–182 (1991).

47. Buonomano, D. V. & Maass, W. State-dependent computations: spatiotemporal processing in cortical networks. Nat Rev Neurosci 10, 113–125 (2009).

48. Ratzlaff, E. H. & Grinvald, A. A tandem-lens epifluorescence macroscope: Hundred-fold brightness advantage for wide-field imaging. Journal of Neuroscience Methods 36, 127–137 (1991).

49. Couto, J. et al. Chronic, cortex-wide imaging of specific cell populations during behavior. Nat Protoc 16, 3241–3263 (2021).

50. Yu, C.-H., Stirman, J. N., Yu, Y., Hira, R. & Smith, S. L. Diesel2p mesoscope with dual independent scan engines for flexible capture of dynamics in distributed neural circuitry. Nat Commun 12, 6639 (2021).

51. Stirman, J. N., Smith, I. T., Kudenov, M. W. & Smith, S. L. Wide field-of-view, multi-region, two-photon imaging of neuronal activity in the mammalian brain. Nat Biotechnol 34, 857–862 (2016).

52. Andryushchenko, E. V. & Guzhov, V. I. Obtaining High-resolution Images with a Large Field of View in Optical Microscopy. in 2024 IEEE 25th International Conference of Young Professionals in Electron Devices and Materials (EDM) 2000–2003 (IEEE, Altai, Russian Federation, 2024). doi:10.1109/EDM61683.2024.10615010.

53. Louth, E. L., Sutton, C. D., Mendell, A. L., MacLusky, N. J. & Bailey, C. D. C. Imaging Neurons within Thick Brain Sections Using the Golgi-Cox Method. JoVE 55358 (2017) doi:10.3791/55358.

54. Stringer, C. & Pachitariu, M. Computational processing of neural recordings from calcium imaging data. Current Opinion in Neurobiology 55, 22–31 (2019).

55. Giovannucci, A. et al. OnACID: Online Analysis of Calcium Imaging Data in Real Time. in Advances in Neural Information Processing Systems (eds Guyon, I. et al.) vol. 30 (Curran Associates, Inc., 2017).

56. Johnson, R. ZeroMQ in Practice: Definitive Reference for Developers and Engineers. (HiTeX Press, 2025).

57. An, G. H. et al. Charuco Board-Based Omnidirectional Camera Calibration Method. Electronics 7, 421 (2018).

58. Lindeberg, T. Scale Invariant Feature Transform. Scholarpedia 7, 10491 (2012).

59. Lakshmi, K. D. & Vaithiyanathan, V. Image Registration Techniques Based on the Scale Invariant Feature Transform. IETE Technical Review 34, 22–29 (2017).

60. Rublee, E., Rabaud, V., Konolige, K. & Bradski, G. ORB: An efficient alternative to SIFT or SURF. in 2011 International Conference on Computer Vision 2564–2571 (IEEE, Barcelona, Spain, 2011). doi:10.1109/ICCV.2011.6126544.

61. Karami, E., Prasad, S. & Shehata, M. Image Matching Using SIFT, SURF, BRIEF and ORB: Performance Comparison for Distorted Images. Preprint at 10.48550/ARXIV.1710.02726 (2017).

62. Morel, J.-M. & Yu, G. ASIFT: A New Framework for Fully Affine Invariant Image Comparison. SIAM J. Imaging Sci. 2, 438–469 (2009).

63. Codreanu, V. et al. GPU-ASIFT: A fast fully affine-invariant feature extraction algorithm. in 2013 International Conference on High Performance Computing & Simulation (HPCS) 474–481 (IEEE, Helsinki, Finland, 2013). doi:10.1109/HPCSim.2013.6641456.

64. Sansoni, G., Corini, S., Lazzari, S., Rodella, R. & Docchio, F. Three-dimensional imaging based on Gray-code light projection: characterization of the measuring algorithm and development of a measuring system for industrial applications. Appl. Opt. 36, 4463 (1997).

65. Pang, Z. P. et al. Induction of human neuronal cells by defined transcription factors. Nature 476, 220–223 (2011).

66. Aharoni, D. CaLab. GitHub (https://github.com/miniscope/CaLab) (2026).

